# Network Dysconnection of Striatum and Default Mode Network Underpins Temporal Dynamics of Depression in Premanifest Huntington’s Disease

**DOI:** 10.1101/2025.08.04.668558

**Authors:** Tamrin Barta, Leonardo Novelli, Nellie Georgiou-Karistianis, Julie Stout, Samantha M Loi, Yifat Glikmann-Johnston, Adeel Razi

## Abstract

**Background and objectives:** Depression emerges years before motor signs in Huntington”s disease gene expansion carriers (HDGECs), yet how directional connectivity patterns evolve over time and relate to depression trajectories remains unknown. We hypothesized HDGECs with depression history would show accelerated grey matter atrophy and disinhibition of striatal-to-default mode network connections over time.

**Methods:** The study utilised the largest longitudinal study of premanifest HD, TrackOn-HD, including structural and resting-state brain scans. HDGECs were stratified by depression history and followed over three annual assessments. Analyses focused on 8 predefined regions spanning striatal and default mode network regions. Here, we applied spectral dynamic causal modelling to examine effective connectivity, alongside voxel-based morphometry of grey matter atrophy and linear mixed models of depressive symptoms. Spectral DCM employed Bayesian model reduction with free energy thresholds ≥0.95 for strong evidence of connectivity changes.

**Results:** The study included 105 HDGECs (53 females, mean age = 43, mean CAG = 43). Depression-related network dysconnection operated independently of regional grey matter atrophy, with HDGECs exhibiting widespread striatal volume loss but no differential atrophy patterns between depression groups. Larger posterior cingulate cortex volumes predicted increased depression severity specifically in HDGECs with depression history, across both Beck Depression Inventory-II (β = 36.79, 95% CI = 2.33-71.25, *p* =.041) and Hospital Anxiety and Depression Scale-Depression Subscale (β = 27.31, 95% CI = 11.12-43.50, *p* =.002). Spectral dynamic causal models explained 88.6% of variance in depression groups and 90.4% in non-depression groups. Longitudinal effective connectivity analyses revealed distinct dysconnection profiles: HDGECs with depression history showed widespread interhemispheric alterations including progressive inhibitory changes of striatal-DMN circuits and aberrant hippocampal regulatory control, while those without depression exhibited more focal dysconnection. These patterns occurred despite no group differences in grey matter atrophy trajectories over 24 months.

**Discussion:** Depression in HDGECs emerges through network dysconnection that operates independently of regional atrophy. Functional alterations may precede detectable structural breakdown and represent pathological responses to subclinical neurodegenerative processes, suggesting functional connectivity markers may serve as early indicators of depression vulnerability in premanifest HD.

## Introduction

Depression emerges early for Huntington”s disease (HD) gene expansion carriers (HDGECs)—years before motor signs signifying manifest diagnosis—with psychiatric disturbance preceding motor dysfunction by up to a decade^1^. Depression symptoms are most severe for HDGECs approaching motor diagnosis^2,3^ and remains elevated throughout manifest HD^4,5^. Within the brain, depression for HDGECs has been linked to structural and functional dysconnection between default mode network (DMN) and striatum^6–8^, however, how network dysconnectivity develops in conjunction with neural atrophy across premanifest HD is not well characterised.

The dorsal striatum—caudate and putamen—degenerates early and severely for HDGECs^9,10^, and forms a critical component in frontostriatal circuitry that shows consistent abnormalities in major depression, including volumetric reductions^11^. In HD, volumetry outside of the striatum meaningfully changes only 20 years after motor onset^12^, suggesting dysconnectivity may be more impactful in the pathophysiology of depression in premanifest HD. For example, frontal region hyperconnectivity is shown over 20 years from manifest diagnosis^13^ becoming hypoconnective with increased disease burden^13,14^. In addition, posterior cingulate cortex (PCC) is proposed as a disease epicentre for HDGECs, with significant structural connectivity reductions closer to motor onset^13^. Both these regions form hubs of the default mode network (DMN). The DMN is as a robust neural marker for major depression^15^, including increased effective connectivity from anterior (including medial prefrontal cortex [MPFC]) to posterior regions (including PCC)^16^. This suggests that the pathogenesis of HD confers depression vulnerability for HDGECs via striatal and DMN dysconnection.

There is conflicting evidence regarding brain change in striatum and DMN and depression in HDGECs. In terms of grey matter atrophy, whole brain or regional volume loss is not associated with broad affective changes^17^, and basal ganglia atrophy is not associated with depressive symptoms^18^. By contrast, depression trajectories—across premanifest and manifest HD—are associated with reduced cortical thickness in limbic regions, prefrontal cortex, and PCC^7^. Looking at striatal and DMN dysconnectivity, there is evidence that depression is associated with increased functional connectivity within DMN regions (but not striatum) and decreased structural connectivity between DMN and basal ganglia^8^. Additionally, there is increased inhibitory striatal influence on DMN (including left putamen), a propensity for right hippocampal involvement, and disinhibitory PCC-hippocampal connectivity, shown in cross-sectional effective connectivity analyses of premanifest HGECs with depression history^6^. These findings suggest posterior DMN as a driver of depression for HDGECs, and potential pathological mechanisms of right DMN. How these directional connectivity patterns evolve over time remains uncharacterised.

Here, we leverage the largest longitudinal dataset (three annual assessments) of HDGECs including structural and functional MRI, TrackOn-HD^10,19^, to investigate grey matter volumetry and effective connectivity aberrations associated with depression. We apply a longitudinal spectral dynamic causal modelling (spDCM)^20– 22^ leveraging Bayesian modelling procedures to discriminate inhibition and excitation of bottom-up and top-down connections, allowing for inference of connectivity direction and strength and within region self-connectivity (i.e., synaptic activity)^23^ between DMN and striatum for premanifest HDGECs with and without a history of depression over 24 months. Further, the framework quantified associations between effective connectivity changes and depressive symptoms.

We hypothesise accelerated grey matter atrophy in regional hubs of left putamen, PCC, hippocampus in HDGECs who have a history of depression. We expect baseline DMN grey matter volume will be associated with depression trajectories, with greater grey matter atrophy associated with increased depression severity. Further, we hypothesise disinhibition of striatum to DMN hubs and decreased regulatory control from DMN to striatum over time for HDGECs. Finally, we expect increasingly disinhibited efferents of regional hubs (right hippocampus, left putamen, PCC) as compensatory mechanisms fail with disease progression.

## Materials and methods

### Participants

This study used Track-On HD data, with detailed recruitment procedures and protocol previously reported^10,19^. Participants gave written informed consent in accordance with the Declaration of Helsinki and procedures were approved by local ethics committees at all sites. All participants were genetically confirmed to have the HD-causing cytosine, adenine, guanine (CAG) expansion (≥ 39 repeats) but did not meet criteria for manifest diagnosis, using the Unified Huntington”s Disease Rating Scale (UHDRS). Exclusion criteria included history of significant head injury and major neurological disorders (excluding HD) at enrolment^10,19^. HDGECs were not excluded based on medication use, unless part of a therapeutic trial, and were not titrated off medications for study sessions. At each timepoint, comorbid depression and associated International Statistical Classification of Diseases and Related Health Problems 10th Revision (ICD10) classifications were collected and verified by trained clinicians. HDGECs were categorized into two groups based on history of depression (current and remitted) or no history of diagnosed depression.

### Clinical measures

Three HD-related variables were included. The CAG repeat length was recorded for each HDGEC. The CAG-Age Product (CAP) score quantified the effect of age and CAG repeat interaction on disease course, equalling 100 at expected diagnosis age^24^. As a measure of disease burden, the HD Integrated Staging System (HD-ISS) was used^25^. The HD-ISS incorporates pathophysiology alongside clinical and functional changes, extending beyond motor impairment by staging disease from 0–3 based on genetic confirmation, striatal volume changes, cognitive impairment, and functional decline^25^.

Regular (daily) mood medication use was recorded at each time point, including antipsychotics, benzodiazepines, selective serotonin reuptake inhibitors (SSRIs), and non-SSRI anxiolytics and antidepressants. At their first available session, 30 HDGECs were taking mood medications (12 HDGECs with a history of depression), with 25 (83.3%) taking one medication.

We included self-report measures to quantify depressive symptoms at each time point: the Beck Depression Inventory-II (BDI-II)^26^ and Hospital Anxiety and Depression Scale-Depression subscale (HADS-D)^27^. The BDI-II consists of 21 items with 4-point Likert scaling (maximum score: 63 and higher scores indicate increased depressive symptoms), measured over a two-week period. The BDI-II has been validated in HD with a cut-off of 10/11 to discriminate clinically elevated depressive symptoms with excellent sensitivity (1.00) and good specificity (0.66)^28^. The HADS-D employs a 4-point Likert scale (maximum score: 21, with higher scores indicating more depressive symptoms)^27^. A cut-off of 6/7 is recommended in HD for discriminating clinically elevated depression, demonstrating excellent sensitivity (1.00) with good specificity (0.82)^28^. We used these thresholds to categorize HDGECs as endorsing clinically elevated depression symptoms at each time-point.

Demographic variables were compared based on depression history using Pearson”s Chi-squared test or ANOVA. Assumptions testing is reported in eResults (eFigure 2-5); minor violations of non-normality were considered acceptable as violations did not impact main analyses.

**Fig. 1.**
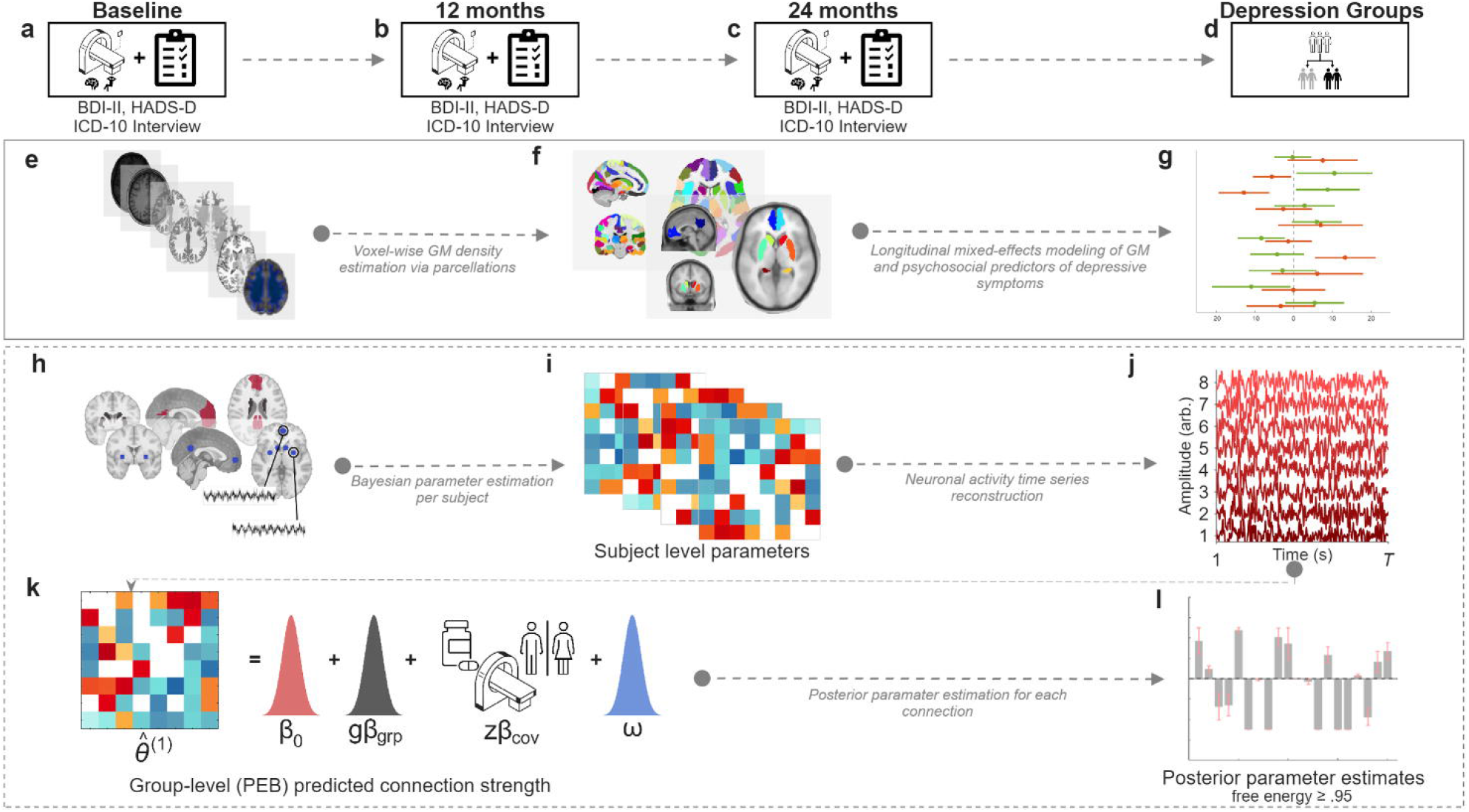
Pipeline for structural volumetry and effective connectivity analyses. **(a-c)** Study protocol encompassing multi-timepoint neuroimaging acquisition at baseline, 12-month, and 24-month intervals, incorporating 3T resting-state functional MRI, structural T1, and comprehensive clinical assessment using Beck Depression Inventory, 2nd Edition (BDI-II), Hospital Anxiety and Depression Scale, Depression subscale (HADS-D), and ICD-10 depression history categorization. **(d)** Participant stratification based on depression history status, enabling group-specific connectivity modelling approaches. **(e-g)** Voxel-based morphometry methods where **(e)** T1 preprocessing pipeline incorporating bias field correction, spatial normalization template space, and tissue segmentation for grey matter volume extraction. (f) Voxel-based morphometry analysis framework demonstrating grey matter probability maps across anatomically defined regions of interest, with color-coded parcellation scheme overlaid on standardized brain template. **(g)** Linear mixed-effects model (LMM) statistical framework examining predictors of depression severity over 24-months, including longitudinal grey matter volume changes. (h-l) Spectral dynamic causal modeling (DCM) preprocessing workflow incorporating **(h)** anatomical region-of-interest definition, blood-oxygen-dependent (BOLD) time-series extraction from eight network nodes spanning default mode network and striatal structures, with spherical masks centered on Montreal Neurological Institute coordinates. **(i)** Subject-specific effective connectivity parameters capture individual deviations from group-level connectivity patterns. **(j)** At the individual level, the DCM forward model generates predicted BOLD time series from effective connectivity parameters, demonstrating the spectral basis for connectivity strength quantification across the 0.1-0.25 Hz frequency range relevant for resting-state network dynamics. **(k)** Shows the predicted connection strength for each region-to-region effect (in Hertz), where θ _i_^(1)^ is participant *i*′s posterior estimate of the connection, β_0_ is the group mean at mean-centered covariates, gβ_grp_ is the group difference term, Zβ_cov_ represents covariate effects (sex, medication, and scanner differences), and ω is the parameter-level residual. **(l)** Posterior parameter estimates, with posterior probability ≥.95, following Bayesian model reduction to identify connections showing reliable group differences.

**Fig. 2.**
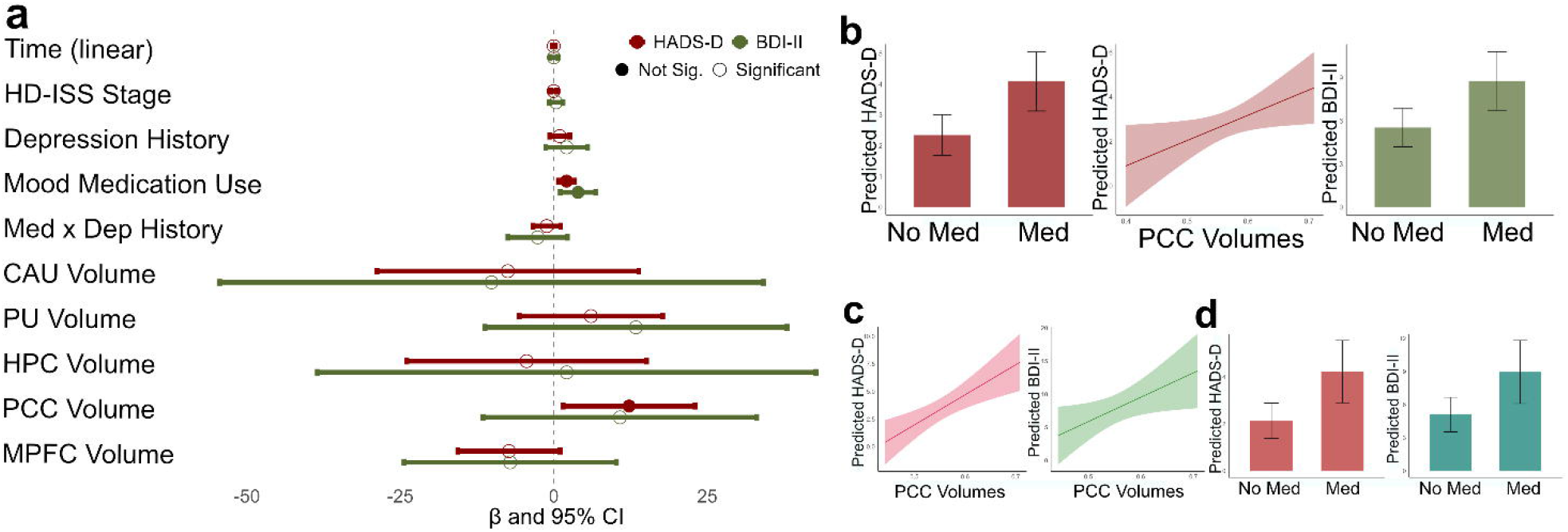
Predictors of Depression Symptoms at 24-month Follow-up. **(a)** Forest plot displaying standardized β coefficients and 95% confidence intervals from linear mixed-effects models predicting Beck Depression Inventory-II (BDI-II; green) and Hospital Anxiety and Depression Scale-Depression subscale (HADS-D; red). Fixed effects include clinical variables (HD-ISS stage, depression history, antidepressant use). All brain volumes are normalized by total intracranial volume (TIV) and scaled (×100) (caudate, putamen, hippocampus, posterior cingulate cortex [PCC], medial prefrontal cortex [MPFC]). Filled circles indicate statistical significance. **(b)** Predicted BDI-II and HADS-D scores stratified by medication use and scatter plot illustrating the relationship between PCC volume and HADS-D scores with regression line and 95% confidence interval (shaded region). **(c)** Scatter plot illustrating the relationship between PCC volume and HADS-D and BDI-II scores with regression line and 95% confidence interval (shaded region), for HDGECs with a history of depression. **(d)** Predicted HADS-D and BDI-II scores stratified by medication class for HDGECs with no depression history.

### Statistical analyses

See Fig. 1 for an outline of the modelling pipeline, described in detail below.

### MRI data acquisition and pre-processing

3T structural (T1) and resting-state functional MRI (fMRI) data over three annual sessions (baseline, 12-month, and 24-month follow-up) were acquired on two different scanner types at four sites: Philips Achieva (Vancouver and Leiden) and Siemens TIM Trio (London and Paris). For T1-weighted image acquisition, a 3D MPRAGE sequence was used with the following parameters: TR 2200ms (Siemens)/ 7.7ms (Philips), TE 2.2ms (S)/3.5ms (P), FOV 280 mm (S)/240 mm (P), flip angle 10°(S)/8°(P), 208(S)/164(P) sagittal slices (slice thickness: 1.1 mm, gap: no gap, matrix size 256 × 256 (S)/224 × 224 (P)) and bandwidth of 240 Hz (S)/241 Hz (P) per participant ^10^. For whole-brain volume acquisition, a T2-weighted echo planar imaging sequence was used with the following parameters: TR 3000 ms, TE 30 ms, FOV 212 mm, flip angle 80°, 48 slices in ascending order (slice thickness: 2.8 mm, gap: 1.5 mm, in plane resolution 3 × 3 × 3 mm) and bandwidth of 1906 Hz per participant.

### MRI preprocessing

Resting-state fMRI preprocessing and quality assurance was conducted using fMRIPrep v21.02.2^29,30^ and MRIQC v22.0.6^31^. The pipeline included slice-timing correction, realignment, spatial normalization to Montreal Neurological Institute (MNI) space, and spatial smoothing with a 6 mm full-width half-maximum (FWHM) Gaussian kernel (eMethods S1.2). A generalized linear model was used to regress white matter and cerebrospinal fluid signals and 6 head motion parameters (3 translation and 3 rotational). ROIs were spherical (radius 6–8 mm) centered on MNI coordinates from prior literature and bounded within masks (eMethods S1.3). Structural T1 preprocessing was conducted using the Computational Anatomy Toolbox (CAT12)^32^, including denoising, bias field correction, and segmentation. Resulting tissue maps were normalized to MNI space using diffeomorphic anatomical registration (DARTEL)^33^ and modulated to preserve absolute tissue volumes, accounting for inter-individual brain size differences. Modulated GM maps were smoothed with a 6mm FWHM Gaussian kernel, in alignment with fMRI preprocessing. Quality assurance procedures in eMethods S1.4.

### Voxel-based morphometry

Voxel-based morphometry (VBM) was implemented using SPM12 and MarsBaR^34^. A flexible factorial design specified subject as a random factor, with time and group as fixed factors, as well as time by group interaction, to characterize volumetric trajectories. Covariates included total intracranial volume (TIV), HD-ISS, and mood medication use. ROI comparisons were conducted using f and t contrasts, and statistical significance (*p* < 0.05) was applied with false discovery rate correction for multiple comparisons across ROIs. Regions were defined using the Tian 3T S1 atlas for striatum and hippocampi^35^ and Schaefer”s 17 network 100 parcellation atlas for MPFC and PCC^36,37^.

### Linear mixed models

Two linear mixed models (LLMs), conducted in R version 4.1.3^38^, examined depressive symptoms, using either HADS-D or BDI-II score as outcome variable over 24-months. Fixed effects were history of depression, TIV-normalized ROI volumes (scaled by 100), mood medication use, and HD-ISS score. Random intercepts accounted for the hierarchical structure of repeated measures nested within participants. P value considered significant at *p* ≤ 0.05 (Analysis pipeline in eMethods S1.5 and model assumptions eFigure 6-7).

### Spectral dynamic causal modelling

spDCM employs a linear state-space model to fit the cross-spectral densities of observed BOLD signals^21^. At the first level, for each participant, fully connected DCM models without exogenous input (i.e., resting state) were specified using our 8 predefined ROIs. Models were subsequently estimated by fitting the DCM forward model (i.e., its parameters) to match the observed cross-spectral densities, yielding estimates of effective connectivity strength between ROI pairs and the corresponding parameter uncertainty.

The longitudinal analysis framework examined connectivity patterns baseline and 24-month follow-up, with two change analyses, one for HDGECS with a depression history and one for HDGECs with no depression history. This approach allowed investigation of connectivity changes associated with depression symptoms within depression subgroups. Covariates included mood medication use and HD-ISS stage to control for confounding effects that varied across timepoints within HDGECs.

The inferred effective connectivity from the first-level analysis was used for hypothesis testing of effects of time within group (i.e., effective connectivity at 24-months with reference to baseline). The Parametric Empirical Bayes (PEB) framework was used to estimate the effect of a history of depression and time for each connection^39^. PEB incorporated both the expected connectivity strength and variance while accounting for between-subject variability and noise. Lastly, Bayesian model reduction was employed as an efficient form of Bayesian model selection^39^. We focused on connections with free energy ≥ 0.95, amounting to strong evidence. Only connections exceeding the threshold of free energy ≥ 0.95 (strong evidence) are reported throughout. Self-connections (inhibitory inputs that prevent runaway positive feedback) modulate excitation-inhibition through synaptic decay regulation^23^, and were analysed alongside inter-regional connectivity parameters. More information is in eMethods S1.6.

## Data availability

No new data were collected for this article. The data that support the findings of this study are available on the Enroll-HD platform via request at https://www.enroll-hd.org/for-researchers/access-data-biosamples/.

## Results

### Participant characteristics

Overall, 105 unique HDGECs were included across analyses. Due to data requirement differences across our VBM, LMM and DCM analyses, as well as quality control for structural and fMRI data, analyses included different participant subsets (Table 1).

**Table 1.** Demographic, clinical, and psychiatric characteristics of HDGECs.

|  | No history of depression (*N* = 75) | History of depression (*N* = 30) | *p* value |
| --- | --- | --- | --- |
| **Demographic Variables** |  |  |  |
| Age |  |  | .983<sup>b</sup> |
| Mean (*sd*) | 43 (9.91) | 43.1 (7.43) |  |
| Range | 23.00 - 68.00 | 28.00 - 60.00 |  |
| Sex |  |  | .422<sup>a</sup> |
| Female | 36 (48.00%) | 17 (56.67%) |  |
| Male | 39 (52.00%) | 13 (43.33%) |  |
| Ethnicity |  |  | .646<sup>a</sup> |
| White | 71 (94.67%) | 29 (96.67%) |  |
| American - Latin | 1 (1.33%) | 1 (3.33%) |  |
| Asian - east | 2 (2.67%) | 0 (0.00%) |  |
| Mixed | 1 (1.33%) | 0 (0.00%) |  |
| Handedness |  |  | .397<sup>a</sup> |
| Right-handed | 69 (92.00%) | 25 (83.33%) |  |
| Left-handed | 3 (4.00%) | 3 (10.00%) |  |
| Ambidextrous | 3 (4.00%) | 2 (6.67%) |  |
| **HD-related Variables** |  |  |  |
| CAG repeat length |  |  | .963<sup>b</sup> |
| Mean (*sd*) | 43.32 (2.38) | 43.1 (1.81) |  |
| Range | 40.00 - 49.00 | 39.00 - 47.00 |  |
| DBS score |  |  | .322<sup>b</sup> |
| Mean (*sd*) | 305.02 (52.68) | 314.18 (55.36) |  |
| Range | 185.00 - 443.00 | 179.00 - 401.00 |  |
| HD-ISS stage |  |  | .476<sup>a</sup> |
| 0 | 23 (30.67%) | 6 (20.00%) |  |
| 1 | 30 (40.00%) | 14 (46.67%) |  |
| 2 | 19 (25.33%) | 7 (23.33%) |  |
| 3 | 3 (4.00%) | 3 (10.00%) |  |
| **Depression variables** |  |  |  |
| Current depression |  |  | < .001<sup>a</sup> |
| No current depression | 75 (100.00%) | 14 (46.67%) |  |
| Current depression | 0 (0.00%) | 16 (53.33%) |  |
| BDI-II score |  |  | .174<sup>b</sup> |
| Mean (*sd*) | 6.01 (7.27) | 8.93 (11.1) |  |
| Range | 0.00 - 27.00 | 0.00 - 46.00 |  |
| HADS-D score |  |  | .063<sup>b</sup> |
| Mean (*sd*) | 2.60 (3.47) | 4.33 (5.2) |  |
| Range | 0.00 - 14.00 | 0.00 - 19.00 |  |
| BDI-II clinical cut-off |  |  | .664<sup>a</sup> |
| Clinically sig. symptoms | 17 (22.67%) | 8 (26.67%) |  |
| Low depressive symptoms | 58 (77.33%) | 22 (73.33%) |  |
| HADS-D clinical cut-off |  |  | .391<sup>a</sup> |
| Clinically sig. symptoms | 10 (13.33%) | 6 (20.00%) |  |
| Low depressive symptoms | 65 (86.67%) | 24 (80.00%) |  |
| Suicidal ideation history |  |  | .073<sup>a</sup> |
| No | 62 (82.67%) | 20 (66.67%) |  |
| Yes | 13 (17.33%) | 10 (33.33%) |  |
| Suicide attempt history |  |  | .231<sup>a</sup> |
| No | 72 (96.00%) | 27 (90.00%) |  |
| Yes | 3 (4.00%) | 3 (10.00%) |  |
| Mood medication use |  |  | < .001<sup>a</sup> |
| No | 15 (45.45%) | 60 (83.33%) |  |
| Yes | 18 (54.55%) | 12 (16.67%) |  |
| Session completed |  |  | - |
| Visit 1 only | 10 (13.33%) | 7 (23.33%) |  |
| Visits 1 & 2 only | 10 (13.33%) | 5 (16.67%) |  |
| Visits 1 & 3 only | 3 (4.00%) | 2 (6.67%) |  |
| Visits 2 & 3 only | 1 (1.33%) | 1 (3.33%) |  |
| All three sessions | 51 (68.00%) | 15 (50.00%) |  |
| **Analysis** |  |  | **Total Obs.** |
| **Voxel-based morphometry** | 53 | 18 | 142 |
| **Linear mixed model** | 61 | 26 | 243 |
| Spectral dynamic causal modelling | 71 | 31 | 164 |

SD = standard deviation. CAG = cytosine-adenine-guanine. CAP = CAG-Age Product. DBS = Disease Burden Score. BDI-II = the Beck Depression Inventory, 2nd Edition. HADS-D = the Hospital Anxiety and Depression Scale, Depression Subscale. Obs = Observations.
<sup>a</sup>Pearson's Chi-squared test
<sup>b</sup>Linear ANOVA

For analyses of brain changes (volumetry and effective connectivity), we used baseline and 24-month follow-up data to capture meaningful brain changes, as cortical atrophy is more subtle in HDGECs^10,40^. At each participant”s first available session, those with a history of depression had significantly higher rates of mood medication use. No other variables differed significantly (Table 1). Depression classifications, episode length and number of episodes are reported in eTable 1. We included regular mood medication use in all analyses. Medications taken, duration, dose, indication, and regime are reported in eTable 1.

### Voxel-based morphometry results

VBM included 71 HDGECs (eTable 3). Patterns of grey matter volume reductions were compared for HDGECs with and without depression history over 24 months (Table 2).

**Table 2.** Grey matter volume changes over 24 months.

| Region | Overall Model |  | Effect of Time |  | No Depression |  | Depression |  | Group × Time |  |
| --- | --- | --- | --- | --- | --- | --- | --- | --- | --- | --- |
|  | F | p<sub>corr</sub> | t | p<sub>corr</sub> | t | p<sub>corr</sub> | t | p<sub>corr</sub> | t | p<sub>corr</sub> |
| CAU<sub>R</sub> | 16.09 | < .001 | 4.34 | < .001 | 4.11 | < .001 | 3.17 | < .001 | 0.26 | .983 |
| CAU<sub>L</sub> | 54.68 | < .001 | 7.85 | < .001 | 8.90 | < .001 | 4.71 | < .001 | -1.56 | 1.000 |
| PU<sub>R</sub> | 16.69 | < .001 | 4.30 | < .001 | 5.65 | < .001 | 2.03 | .172 | -1.94 | 1.000 |
| PU<sub>L</sub> | 31.95 | < .001 | 6.97 | < .001 | 7.20 | < .001 | 4.67 | < .001 | -0.42 | 1.000 |
| HPC<sub>R</sub> | 6.79 | < .001 | 0.24 | 1.000 | 0.77 | .937 | -0.21 | .985 | -0.74 | 1.000 |
| HPC<sub>L</sub> | 7.94 | < .001 | 2.19 | .164 | 2.04 | .181 | 1.63 | .232 | 0.18 | .989 |
| PCC | 24.78 | < .001 | -1.06 | .946 | -1.13 | 1.000 | -0.69 | 1.000 | 0.11 | .992 |
| MPFC | 14.18 | < .001 | 0.30 | 1.000 | 1.50 | .327 | -0.63 | 1.000 | -1.66 | 1.000 |

p<sub>corr</sub> = p-values FWE-corrected at the ROI level. Group × Time Interaction tests whether volumetric changes from Time 1 to Time 2 were more pronounced in HDGECs depression history. MPFC = Medial Prefrontal Cortex; PCC = Posterior Cingulate Cortex; HPC<sub>L</sub> = Left Hippocampus; HPC<sub>R</sub> = Right Hippocampus; CAU<sub>L</sub> = Left Caudate; PU<sub>L</sub> = Left Putamen; PU<sub>R</sub> = Right Putamen.

**Table 3.**
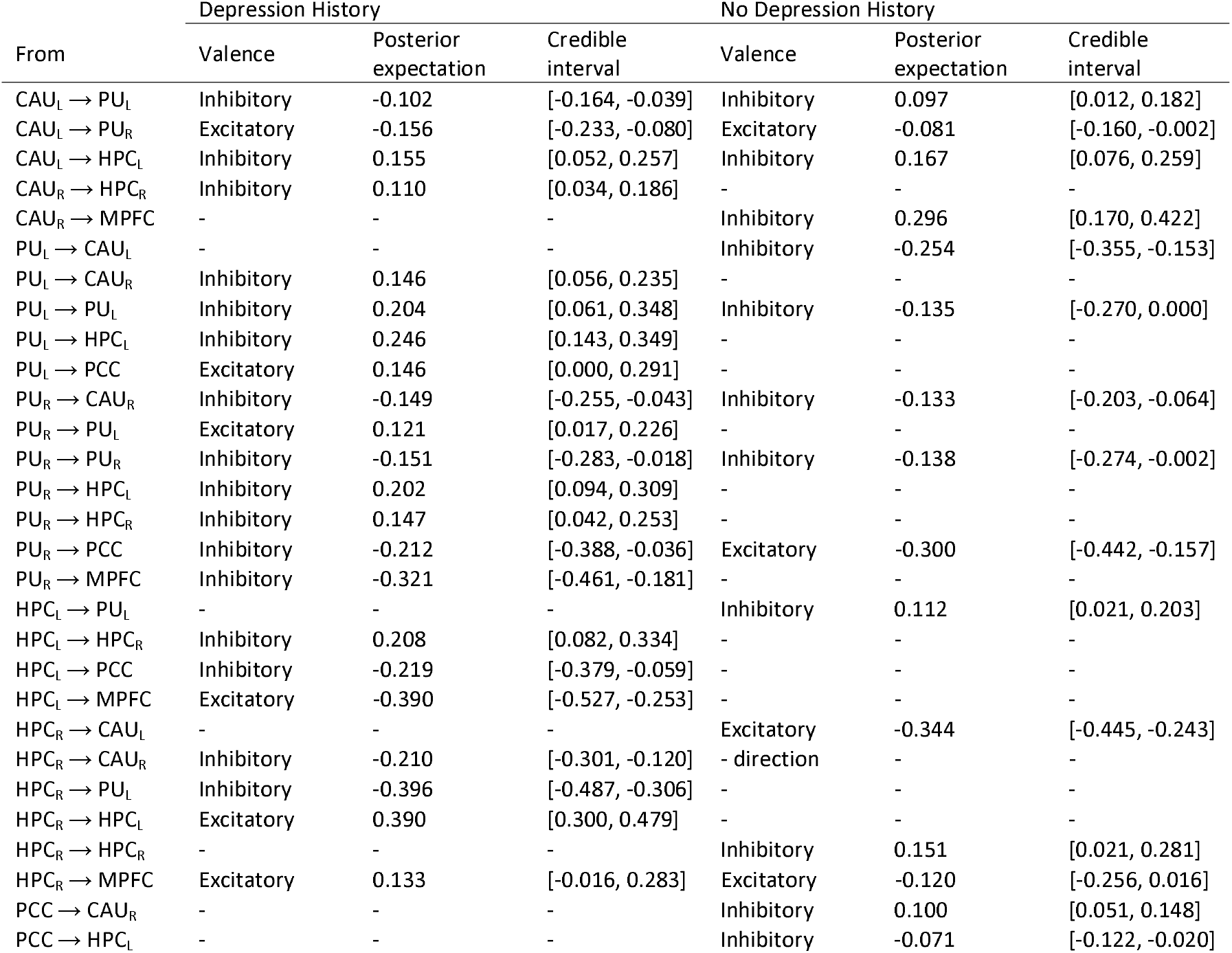
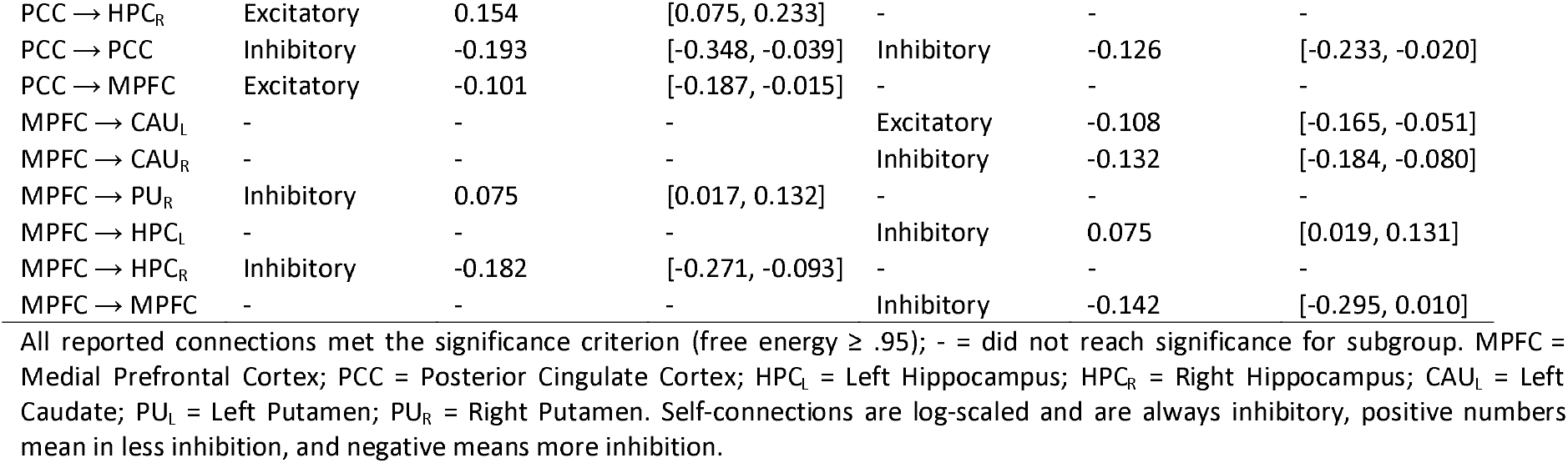
Between-region effective connectivity changes for HDGECs by depression history status.

| From | Depression History |  |  | No Depression History |  |  |
| --- | --- | --- | --- | --- | --- | --- |
|  | Valence | Posterior expectation | Credible interval | Valence | Posterior expectation | Credible interval |
| CAU<sub>L</sub> → PU<sub>L</sub> | Inhibitory | -0.102 | [-0.164, -0.039] | Inhibitory | 0.097 | [0.012, 0.182] |
| CAU<sub>L</sub> → PU<sub>R</sub> | Excitatory | -0.156 | [-0.233, -0.080] | Excitatory | -0.081 | [-0.160, -0.002] |
| CAU<sub>L</sub> → HPC<sub>L</sub> | Inhibitory | 0.155 | [0.052, 0.257] | Inhibitory | 0.167 | [0.076, 0.259] |
| CAU<sub>R</sub> → HPC<sub>R</sub> | Inhibitory | 0.110 | [0.034, 0.186] | - | - | - |
| CAU<sub>R</sub> → MPFC | - | - | - | Inhibitory | 0.296 | [0.170, 0.422] |
| PU<sub>L</sub> → CAU<sub>L</sub> | - | - | - | Inhibitory | -0.254 | [-0.355, -0.153] |
| PU<sub>L</sub> → CAU<sub>R</sub> | Inhibitory | 0.146 | [0.056, 0.235] | - | - | - |
| PU<sub>L</sub> → PU<sub>L</sub> | Inhibitory | 0.204 | [0.061, 0.348] | Inhibitory | -0.135 | [-0.270, 0.000] |
| PU<sub>L</sub> → HPC<sub>L</sub> | Inhibitory | 0.246 | [0.143, 0.349] | - | - | - |
| PU<sub>L</sub> → PCC | Excitatory | 0.146 | [0.000, 0.291] | - | - | - |
| PU<sub>R</sub> → CAU<sub>R</sub> | Inhibitory | -0.149 | [-0.255, -0.043] | Inhibitory | -0.133 | [-0.203, -0.064] |
| PU<sub>R</sub> → PU<sub>L</sub> | Excitatory | 0.121 | [0.017, 0.226] | - | - | - |
| PU<sub>R</sub> → PU<sub>R</sub> | Inhibitory | -0.151 | [-0.283, -0.018] | Inhibitory | -0.138 | [-0.274, -0.002] |
| PU<sub>R</sub> → HPC<sub>L</sub> | Inhibitory | 0.202 | [0.094, 0.309] | - | - | - |
| PU<sub>R</sub> → HPC<sub>R</sub> | Inhibitory | 0.147 | [0.042, 0.253] | - | - | - |
| PU<sub>R</sub> → PCC | Inhibitory | -0.212 | [-0.388, -0.036] | Excitatory | -0.300 | [-0.442, -0.157] |
| PU<sub>R</sub> → MPFC | Inhibitory | -0.321 | [-0.461, -0.181] | - | - | - |
| HPC<sub>L</sub> → PU<sub>L</sub> | - | - | - | Inhibitory | 0.112 | [0.021, 0.203] |
| HPC<sub>L</sub> → HPC<sub>R</sub> | Inhibitory | 0.208 | [0.082, 0.334] | - | - | - |
| HPC<sub>L</sub> → PCC | Inhibitory | -0.219 | [-0.379, -0.059] | - | - | - |
| HPC<sub>L</sub> → MPFC | Excitatory | -0.390 | [-0.527, -0.253] | - | - | - |
| HPC<sub>R</sub> → CAU<sub>L</sub> | - | - | - | Excitatory | -0.344 | [-0.445, -0.243] |
| HPC<sub>R</sub> → CAU<sub>R</sub> | Inhibitory | -0.210 | [-0.301, -0.120] | - direction | - | - |
| HPC<sub>R</sub> → PU<sub>L</sub> | Inhibitory | -0.396 | [-0.487, -0.306] | - | - | - |
| HPC<sub>R</sub> → HPC<sub>L</sub> | Excitatory | 0.390 | [0.300, 0.479] | - | - | - |
| HPC<sub>R</sub> → HPC<sub>R</sub> | - | - | - | Inhibitory | 0.151 | [0.021, 0.281] |
| HPC<sub>R</sub> → MPFC | Excitatory | 0.133 | [-0.016, 0.283] | Excitatory | -0.120 | [-0.256, 0.016] |
| PCC → CAU<sub>R</sub> | - | - | - | Inhibitory | 0.100 | [0.051, 0.148] |
| PCC → HPC<sub>L</sub> | - | - | - | Inhibitory | -0.071 | [-0.122, -0.020] |
| PCC → HPC<sub>R</sub> | Excitatory | 0.154 | [0.075, 0.233] | - | - | - |
| PCC → PCC | Inhibitory | -0.193 | [-0.348, -0.039] | Inhibitory | -0.126 | [-0.233, -0.020] |
| PCC → MPFC | Excitatory | -0.101 | [-0.187, -0.015] | - | - | - |
| MPFC → CAU<sub>L</sub> | - | - | - | Excitatory | -0.108 | [-0.165, -0.051] |
| MPFC → CAU<sub>R</sub> | - | - | - | Inhibitory | -0.132 | [-0.184, -0.080] |
| MPFC → PU<sub>R</sub> | Inhibitory | 0.075 | [0.017, 0.132] | - | - | - |
| MPFC → HPC<sub>L</sub> | - | - | - | Inhibitory | 0.075 | [0.019, 0.131] |
| MPFC → HPC<sub>R</sub> | Inhibitory | -0.182 | [-0.271, -0.093] | - | - | - |
| MPFC → MPFC | - | - | - | Inhibitory | -0.142 | [-0.295, 0.010] |

All reported connections met the significance criterion (free energy $\geq .95$ ); - = did not reach significance for subgroup. MPFC = Medial Prefrontal Cortex; PCC = Posterior Cingulate Cortex; HPC<sub>L</sub> = Left Hippocampus; HPC<sub>R</sub> = Right Hippocampus; CAU<sub>L</sub> = Left Caudate; PU<sub>L</sub> = Left Putamen; PU<sub>R</sub> = Right Putamen. Self-connections are log-scaled and are always inhibitory, positive numbers mean in less inhibition, and negative means more inhibition.

When looking at all HDGECs and for HDGECs with no history of depression, all striatal regions exhibited significant volume loss, however, there was more selective striatal volume loss in HDGECs with a history of depression, with no significant reduction in right putamen grey matter. No DMN regions showed significant volume changes over 24 months in either group. Group by time interactions revealed no differential atrophy patterns between HDGECs with and without depression history across any examined region.

### Linear mixed model of depression symptoms

LMMs included 87 HDGECs (eTable 4). LLMs with linear time trajectories demonstrated optimal fit across both models (see eResults). There was substantial between-subject heterogeneity and ICC indicated individual differences accounted for 73.2% and 68.5% of total variance in HADS-D and BDI-II scores, respectively. Variance explained by fixed effects alone was modest (marginal R^2^: HADS-D = 0.119; BDI-II = 0.072), with random intercepts increasing explanatory power (conditional R^2^: HADS-D = 0.764; BDI-II = 0.708). Fig. 2 shows the results as a forest plot of standardized β coefficients and 95% confidence intervals (eTable 5).

### Fixed effects predicting depressive symptoms

Increased mood medication use was associated with both elevated BDI-II (*p* =.008) and HADS-D scores (*p* =. 003; Fig. 2a). There was significant and overlapping variability in predicted scores (shown in error bars of Fig. 2b). Additionally, larger PCC volumes associated with increased depression severity on the HADS-D (*p* =.03), (Fig. 2b).

When analyses were constrained to HDGECs with a history of depression, larger PCC volumes associated with more severe depression on BDI-II, β = 36.79, *SE* = 16.88, *t*(19.70) = 2.18, *p* =.041, and HADS-D, β = 27.31, *SE* = 7.85, *t*(22.04) = 3.48, *p* =.002 (Fig. 2c), however, for HDGECs with no history of depression, anti-depressant usage was associated with increased depression severity on HADS-D, β = 2.10, *SE* = 0.68, *t*(158.43) = 3.08, *p* =.002, and BDI-II, β = 3.86, *SE* = 1.49, *t*(150.87) = 2.60, *p* =.010 (Fig. 2d). See eFigure 8-9 for forest plots of standardized β coefficients.

### Effective Connectivity Changes

#### DMN and striatal effective connectivity changes over 24-months for HDGECS with and without depression history

Analyses included 102 HDGECs (eTable 6). The estimation of DCM models for HDGECs with a history of depression was excellent, with mean variance explained by the model of 88.6% (*SD* = 4.5, range = 74.12-94.31; baseline = 88.5%, 24-month follow-up = 88.7%). Results were similar for HDGECs with no history of depression, with mean variance explained by the model of 90.4% (*SD* = 3.8, range = 75.15-96.45; baseline = 90.5%, 24-month follow-up = 90.3%). Fig. 3 presents the effective connectivity changes observed across HDGECs stratified by depression history over 24-months.

**Fig. 3.**
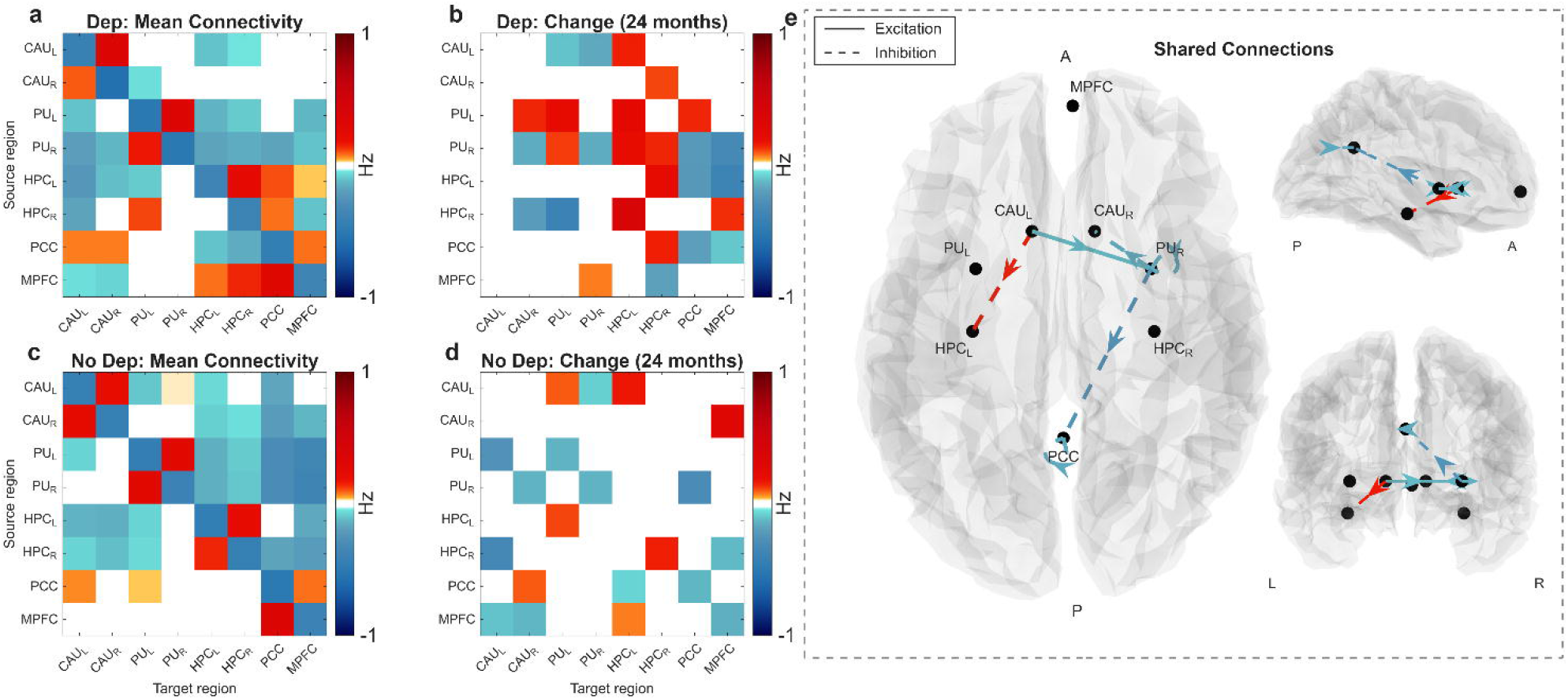
Effective connectivity profiles reveal depression-specific and shared network alterations in HDGECs. Baseline effective connectivity matrix for **(a)** HDGECs with depression history and **(c)** HDGECs without depression history, showing mean connectivity strength across default mode regions (medial prefrontal cortex [MPFC], posterior cingulate cortex [PCC], bilateral hippocampi [HPCL, HPCR]) and striatal structures (bilateral caudate [CAUL, CAUR] and putamen [PUL, PUR]). Color scale represents connection strength in Hz. blue represents inhibitory efferents, while red represents excitatory efferent connections. Longitudinal (24 months) effective connectivity changes for **(c)** HDGECs with depression history and **(d)** HDGECs without depression history. Red indicates increasing connectivity and blue represents decreasing connectivity. Self-connections are log-scaled and are always inhibitory, positive numbers mean in less inhibition, and negative means more inhibition. **(e)** Shared connectivity modifications common to both HDGEC groups, anatomically visualized across axial, coronal, and sagittal brain planes. Solid lines represent excitatory connections while dashed lines indicate inhibitory influences. Red arrows denote strengthening shared connections and blue arrows represent decreasing shared pathways. Anatomical orientation: A = anterior, P = posterior, L = left, R = right. All connections displayed meet 95% posterior probability threshold for statistical significance.

Maximum *a posteriori* estimates showed similar mean effective connectivity profiles for HDGECs with depression history (Fig. 3a) and those without depression history (Fig. 3c), with notable inhibitory influence of striatum on DMN. The two spDCM analyses identified three distinct profiles of efferent connectivity changes: HDGECs with depression history alterations (Fig. 3b), HDGECs with no depression history specific changes (Fig. 3d), and shared connectivity changes (Fig. 3e; plotted on a brain mesh in axial, coronal and sagittal planes). Posterior expectation and connection strength differed for shared connections and are reported in Table 3, alongside including valence and credible intervals.

#### Effective connectivity changes and depressive symptoms

There were no significant differences in depression symptom severity between baseline and 24 months for BDI-II, *t*(60) = -0.074, *p* = 0.94, or HADS-D, *t*(60) = 0.057, *p* = 0.95. Therefore, we used significant connections for each group to see if the change in connectivity over 24 months was associated with clinically elevated depression symptoms at 24-month follow-up, for HDGECs with a history of depression and those without (Fig. 4).

**Fig. 4.**
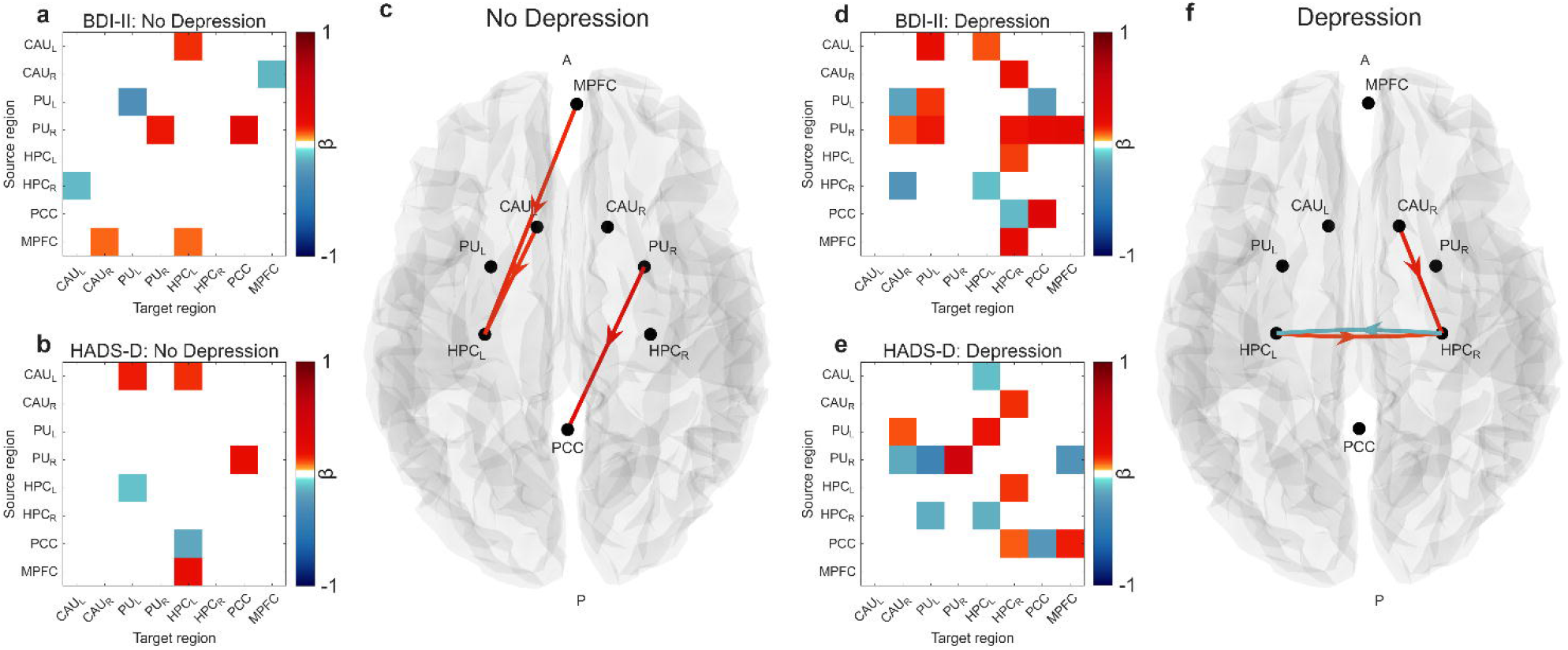
Depression severity associations and predictive connectivity patterns by depression history. Depression severity associations with effective connectivity patterns across Beck Depression Inventory, 2nd Edition [BDI-II] and Hospital Anxiety and Depression Scale, Depression subscale [HADS-D] clinical cut-off measures. **(a)** BDI-II and **(b)** HADS-D clinically elevated depression cut-off associations with effective connectivity for HDGECs without depression history **(d)** BDI-II and **(e)** HADS-D clinically elevated depression cut-off associations with effective connectivity for HDGECs with depression history. Common connectivity associations across depression measures for **(c)** HDGECs without depression history and **(f)** HDGECs with depression history, anatomically visualized on sagittal brain plane. Red connections indicate positive associations with depression severity while blue connections represent negative associations. Connection matrices display directed inter-regional influences between default mode network regions (medial prefrontal cortex [MPFC], posterior cingulate cortex [PCC], bilateral hippocampi [HPCL, HPCR]) and striatal structures (bilateral caudate [CAUL, CAUR] and putamen [PUL, PUR]). Color scale represents standardized connectivity coefficients (β). Anatomical orientation: A = anterior, P = posterior. Positive evidence set at 95% posterior probability threshold for statistical significance.

## Discussion

Here we show that changes in effective connectivity of DMN and striatum differentiate HDGECs with a history of depression from those without. Moreover, these changes occur in the absence of differentiating patterns of grey matter atrophy in any region. Larger PCC volumes correlated with increased depression severity— specifically in HDGECs with a history of depression—contradicting the classical neuroscientific view linking structural preservation to reduced vulnerability. Instead, our findings contribute to a growing literature that suggests functional rearrangement and pathological processes of relatively preserved regions in networks underpinning the phenomenology of HD.

### Grey matter atrophy does not differentiate HDGECs with depression

VBM analyses showed striatal degeneration across all HDGECs, with no difference in atrophy patterns across by depression history status over 24 months. Our findings replicate findings of early and frank atrophy of grey matter in the striatum^9,10^, however, we failed to find a relationship between brain atrophy and depression. In non-neurological populations, people with major depression show grey matter atrophy in prefrontal regions compared to non-depressed controls^41^, however, our findings show depression in premanifest HD is not related to atrophy of depression-related regions. This extends prior research which found no associations between striatal atrophy and depressive symptoms in HDGECs^18^ to include depression-related regions. It is possible that depression does confer risk for accelerated atrophy in later stages, as seen in Parkinson”s disease where depression severity predicts faster cortical thinning rates^42^. It”s possible that similar patterns of atrophy trajectories may appear over longer time periods, including the transition to manifest HD, in our depression-related regions. Our sample comprised HDGECs in early stages of premanifest HD; with 73% of HDGECs falling in HD-ISS stage 0 or 1, representing minimal biological consequences of HD. We may have missed the period of volumetric changes that would be associated with depression in HDGECs. To further explore this relationship, analyses using depression severity to try predict cortical atrophy would help elucidate whether brain atrophy follows similar or divergent patterns over disease progression.

We show that larger PCC volumes associate with depression severity for the HADS-D, regardless of depression history. Critically, when we looked at HDGECs with a history of depression, we saw that this relationship extended to the BDI-II, however, associations were lost for HDGECs without a history of depression. This suggests an alteration relationship between volumetry and depression severity in HDGECs with a comorbid depression history, and may represent functional reorganisation unique to depression in HD. Broadly, the PCC has been proposed as a disease epicenter for HDGECs, with significant structural connectivity reductions^13^, and in major depression, more severe depressive mood has been associated with larger PCC volumes in older adults^43^. The disease process for HDGECs may increase risk for more severe depressive symptoms, and links disparate threads to suggest PCC functional reorganisation as a depression-specific mechanism. Previous findings, based on a single question regarding low mood, found reduced cortical thickness in DMN regions (ventromedial prefrontal cortex, medial temporal gyrus, PCC), as well as reduced grey matter in ventromedial prefrontal cortex and middle temporal gyri, with concurrent increased grey matter in right caudate^7^. One key limitation of previous investigations was collapsing premanifest HDGECs with manifest HD, making it difficult to tease apart potential asynchronous and asymmetric patterns of neurodegeneration and depression. Our findings elucidate that patterns of grey matter reductions may be driven by changes in later HD periods. This temporal sequence suggests a biphasic model of depression-related structural change in HD: initial functional reorganisation in key network hubs during premanifest HD, followed by progressive atrophy as disease advances.

### Effective connectivity changes of depression for HDGECs

Effective connectivity findings extend previous cross-sectional work^6^ while revealing the temporal evolution of network dysfunction. For HDGECs with a history of depression, connectivity changes over 24 months were more widespread and interhemispheric. In keeping with previous findings in major depression^44^ and cross-sectional work^6^, there was a propensity for right hemisphere effective connectivity aberrations. This is in contrast with previous work in HD showing leftward putamen grey matter loss associated with emotion processing more broadly^45^; while left putamen had decreasing inhibitory control on both right caudate and left hippocampus, effective connectivity patterns show bilateral putamen involvement in depression trajectories. When we look at specific patterns, there is disinhibition of MPFC on right putamen and conversely increasing inhibitory control of right putamen on MPFC. In previous cross-sectional work, right putamen had reduced inhibitory control over MPFC^6^, and this appears to become increasingly inhibitory over 24-months. This is in line with hypotheses. In terms of hippocampal connectivity, left hippocampus has inhibitory influence on both hubs of the DMN, while right hippocampus has reduced excitatory influence on the striatum. Cross-sectional work showed right hippocampus inhibitory control over left caudate^6^, suggesting dynamic alterations in effective connectivity over disease progression. These findings together suggest inhibitory changes associated with depression for HDGECs over premanifest HD, which aligns with accounts of hypoconnectivity in HD with disease progression.

Changes in effective connectivity of HDGECs without depression provide insight into dysconnectivity related to HD pathogenesis. We found evidence of aberrant MPFC control over bilateral caudate, that is increasingly inhibitory in left hemisphere and hypoexcitatory in the right. Additionally, HD progression is associated with aberrant self-connectivity of MPFC, suggesting accelerated synaptic decay. Right hippocampus inhibition is influential in HD pathogenesis with reduced synaptic decay (reduced inhibition) and hypoexcitation with left caudate. Interestingly, we saw that at baseline, increasing inhibition of right hippocampus to right caudate differentiate HDGECs with depression history from those without^6^. This suggests that as HD progresses, right hippocampus connectivity may break down and become maladaptive as neurodegenerative burden increases. Some shared connections across groups showed opposite valences. Left putamen self-connectivity and right hippocampus to MPFC connectivity was less inhibitory in HDGECS with depression history compared to more inhibitory in HDGECs with no depression, while left caudate to putamen was increasing inhibitory in HDGECS with depression history compared to reduced inhibition in HDGECs with no depression over 24 months. These findings suggest depression in HDGECs results in aberrant effective connectivity changes in previously proposed epicenters for depression in premanifest HD^6^.

Effective connectivity associations with depression severity reveals distinct neural network signatures that differentiate HDGECs by depression history status. For HDGECs with depression history, both BDI-II and HADS-D severity measures showed extensive connectivity associations spanning default mode network and striatal regions, with prominent involvement of bilateral hippocampal connectivity. In contrast, HDGECs without depression history demonstrated markedly sparse connectivity associations with depression severity measures, primarily involving aberrant top-down control with limited hippocampal efferents. These findings suggest HDGECs without depression history maintain greater neural flexibility to respond to mood-related dysfunction, with depression severity associations confined to discrete top-down circuitry. Conversely, HDGECs with depression history demonstrate more diffuse network engagement, potentially reflecting progressive integration of depression-related network dysfunction with core HD pathophysiology. This pattern suggests depression in HD creates more pervasive vulnerability to mood-related network alterations, where depression severity becomes associated with broader circuit dysfunction.

Our findings motivate several future directions. Volumetric changes represent one dimension of structural brain organization and examine structure and function independently, rather than modelling structural and effective connective coupling. There is evidence that depression for premanifest HDGECs is associated with increased functional connectivity within DMN regions (but not striatum) and decreased structural connectivity between DMN and basal ganglia^8^. The mechanistic relationship between structural and functional connectivity—termed structure-function coupling—has emerged to investigate disease-related network reorganization^46^. A question arises whether neurodegeneration forces functional networks to be constrained by structural dysconnection, or if HD pathogenesis engenders functional-structural decoupling through compensatory mechanisms. Empirical research in Alzheimer”s disease and major depression provides some preliminary evidence with conflicting evidence of progressive structure-function decoupling of whole brain networks^47^ contrasting with increased structure-function coupling DMN in Alzheimer”s disease^48^ and major depression^49^. Changes in the relationship between structure and function appear to be network specific but current investigations remain correlational and non-deterministic. Effective connectivity analyses, including novel models that leverage structural connectivity-based group level priors for DCM^50^, are well placed to elucidate whether depression arises from changes to the structure-function relationship for premanifest HDGECs.

Our findings suggests that depression for HDGECs emerges through network dysconnection that operates independently of regional atrophy. Functional alterations may precede detectable structural breakdown in the DMN and represent early compensatory or pathological responses to subclinical neurodegenerative processes.

## Supplementary material

Supplementary material is available online.

