## Supplementary Materials for "Network Dysconnection of Striatum and Default Mode Network Underpins Temporal Dynamics of Depression in Premanifest Huntington’s Disease"

**Supplementary Materials: Temporal Dynamics of Depression in Huntington's Disease Gene Expansion Carriers: A Network Dysconnection Approach**

Tamrin Barta^1^, Leonardo Novelli^1^, Nellie Georgiou-Karistianis^1^, Julie Stout^1^, Samantha Loi^2,3^, Yifat Glikmann-Johnston^1^, and Adeel Razi^1,4,5,6^

Author affiliations:

1 Turner Institute of Brain and Mental Health at the School of Psychological Sciences, and, Faculty of Medicine, Nursing and Health Sciences, Monash University

2 Department of Psychiatry, University of Melbourne.

3 Neuropsychiatry Centre, Royal Melbourne Hospital, Parkville, Australia

4 Monash Biomedical Imaging, Monash University, Clayton, Victoria, Australia

5 Wellcome Centre for Human Neuroimaging, University College London, London, United Kingdom

6 CIFAR Azrieli Global Scholars Program, CIFAR, Toronto, Ontario, Canada

### S1. eMethods

#### S1.1 VBM Preprocessing

When multiple sessions were available, CAT12's longitudinal pipeline utilized subject-specific templates and minimized morphometric estimation bias across time points^3^. Pre-processing included spatially adaptive non-local means denoising (SANLM)^1^, bias field correction, and unified segmentation^2^. Chosen parcels (eFigure 1) included: MPFC (Schaefer 17 Network 100 Parcel: 40 and 92), PCC (Schaefer 17 Network 100 Parcel: 39 and 91), right hippocampus (Tian Subcortical Atlas S1 Parcel: 1), left hippocampus (Tian Subcortical Atlas S1 Parcel: 9), right putamen (Tian Subcortical Atlas S1 Parcel: 7), left putamen (Tian Subcortical Atlas S1 Parcel: 15), right caudate (Tian Subcortical Atlas S1 Parcel: Parcel 8) and left caudate (Tian Subcortical Atlas S1 Parcel: 16). Total intracranial volumes (TIV) was included as a covariate to account for inter-individual differences in overall brain size^2^.

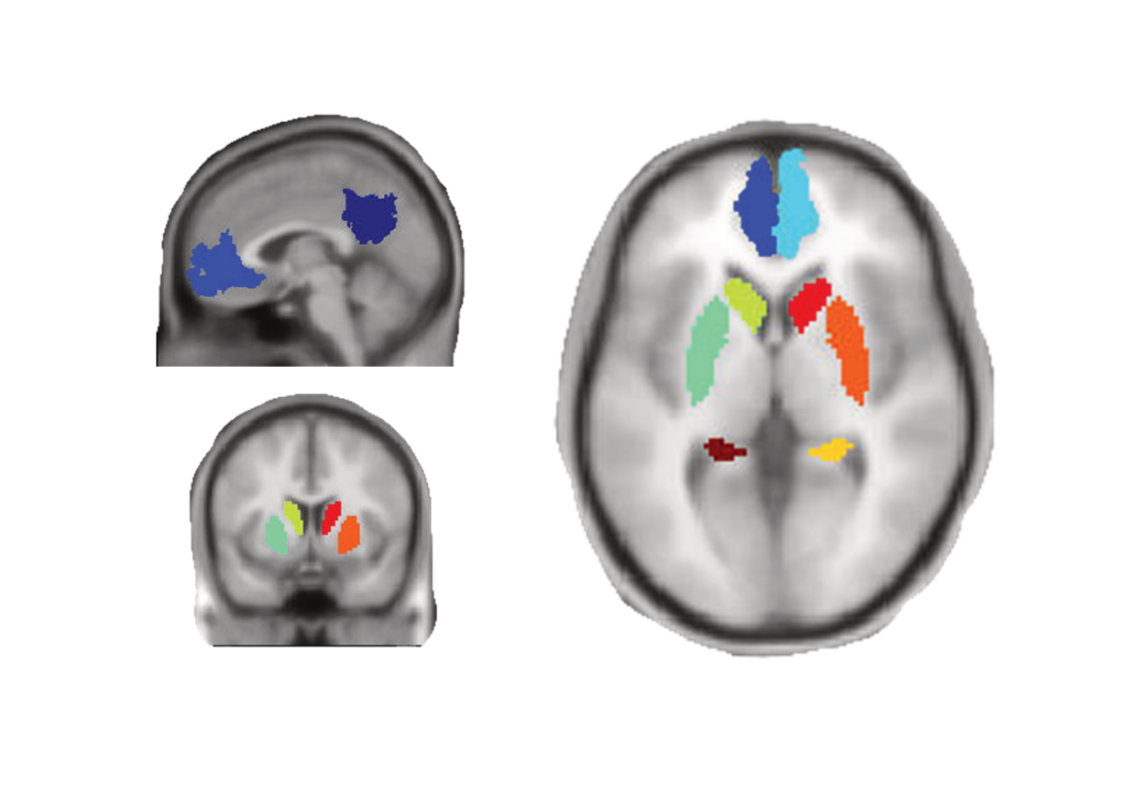

eFigure 1 Final Parcellations for Voxel-Based Morphometry Analyses. (Top Left) Sagittal view. Blue is the medial prefrontal cortex, and navy is the posterior cingulate cortex. (Bottom Left) Coronal view. Green and red show the left and right caudate respectively, and mint and orange show the left and right putamen respectively. (Right) Axial view. Blue and light blue show the left and right medial prefrontal cortex respectively, green and red show the left and right caudate respectively, mint and orange show the left and right putamen respectively, and the brown and yellow show the left and right hippocampus respectively.

#### S1.2 fMRIPrep

Results included in this manuscript come from preprocessing performed using *fMRIPrep* 21.0.2^4,5^ (RRID:SCR_016216), which is based on *Nipype* 1.6.1^6,7^ (RRID:SCR_002502).

##### S1.2.1 Anatomical data preprocessing

A total of 3 T1-weighted (T1w) images were found within the input BIDS dataset. All of them were corrected for intensity non-uniformity (INU) with N4BiasFieldCorrection^8^, distributed with ANTs 2.3.3^9^(RRID:SCR_004757). The T1w-reference was then skull-stripped with a *Nipype* implementation of the antsBrainExtraction.sh workflow (from ANTs), using OASIS30ANTs as target template. Brain tissue segmentation of cerebrospinal fluid (CSF), white-matter (WM) and gray-matter (GM) was performed on the brain-extracted T1w using fast (FSL 6.0.5.1:57b01774, RRID:SCR_002823)^10^. A T1w-reference map was computed after registration of 3 T1w images (after INU-correction) using mri_robust_template (FreeSurfer 6.0.1)^11^. Brain surfaces were reconstructed using recon-all (FreeSurfer 6.0.1, RRID:SCR_001847)^12^, and the brain mask estimated previously was refined with a custom variation of the method to reconcile ANTs-derived and FreeSurfer-derived segmentations of the cortical gray-matter of Mindboggle (RRID:SCR_002438)^13^. Volume-based spatial normalization to one standard space (MNI152NLin2009cAsym) was performed through nonlinear registration with antsRegistration (ANTs 2.3.3), using brain-extracted versions of both T1w reference and the T1w template. The following template was selected for spatial normalization: *ICBM 152 Nonlinear Asymmetrical template version 2009c* [RRID:SCR_008796; TemplateFlow ID: MNI152NLin2009cAsym]^14^.

##### S1.2.2 Functional data preprocessing

For each of the 3 BOLD runs found per subject (across all tasks and sessions), the following preprocessing was performed. First, a reference volume and its skull-stripped version were generated using a custom methodology of *fMRIPrep*. Head-motion parameters with respect to the BOLD reference (transformation matrices, and six corresponding rotation and translation parameters) are estimated before any spatiotemporal filtering using mcflirt (FSL 6.0.5.1:57b01774)^15^. BOLD runs were slice-time corrected to 1.48s (0.5 of slice acquisition range 0s-2.95s) using 3dTshift from AFNI ^16^(RRID:SCR_005927). The BOLD time-series (including slice-timing correction when applied) were resampled onto their original, native space by applying the transforms to correct for head-motion. These resampled BOLD time-series will be referred to as *preprocessed BOLD in original space*, or just *preprocessed BOLD*. The BOLD reference was then co-registered to the T1w reference using bbregister (FreeSurfer) which implements boundary-based registration^17^. Co-registration was configured with six degrees of freedom. Several confounding time-series were calculated based on the *preprocessed BOLD*: framewise displacement (FD), DVARS and three region-wise global signals. FD was computed using two formulations following Power (absolute sum of relative motions^18^, and Jenkinson (relative root mean square displacement between affines ^15^). FD and DVARS are calculated for each functional run, both using their implementations in *Nipype* (following the definitions by Power et al. 2014). The three global signals are extracted within the CSF, the WM, and the whole-brain masks. Additionally, a set of physiological regressors were extracted to allow for component-based noise correction (*CompCor*^19^). Principal components are estimated after high-pass filtering the *preprocessed BOLD* time-series (using a discrete cosine filter with 128s cut-off) for the two *CompCor* variants: temporal (tCompCor) and anatomical (aCompCor). tCompCor components are then calculated from the top 2% variable voxels within the brain mask. For aCompCor, three probabilistic masks (CSF, WM and combined CSF+WM) are generated in anatomical space. The implementation differs from that of Behzadi et al. in that instead of eroding the masks by 2 pixels on BOLD space, the aCompCor masks are subtracted a mask of pixels that likely contain a volume fraction of GM. This mask is obtained by dilating a GM mask extracted from the FreeSurfer’s *aseg* segmentation, and it ensures components are not extracted from voxels containing a minimal fraction of GM. Finally, these masks are resampled into BOLD space and binarized by thresholding at 0.99 (as in the original implementation). Components are also calculated separately within the WM and CSF masks. For each CompCor decomposition, the *k* components with the largest singular values are retained, such that the retained components’ time series are sufficient to explain 50 percent of variance across the nuisance mask (CSF, WM, combined, or temporal). The remaining components are dropped from consideration. The head-motion estimates calculated in the correction step were also placed within the corresponding confounds file. The confound time series derived from head motion estimates and global signals were expanded with the inclusion of temporal derivatives and quadratic terms for each ^20^. Frames that exceeded a threshold of 0.5 mm FD or 1.5 standardised DVARS were annotated as motion outliers. The BOLD time-series were resampled into standard space, generating a *preprocessed BOLD run in MNI152NLin2009cAsym space*. First, a reference volume and its skull-stripped version were generated using a custom methodology of *fMRIPrep*. All resamplings can be performed with *a single interpolation step* by composing all the pertinent transformations (i.e. head-motion transform matrices, susceptibility distortion correction when available, and co-registrations to anatomical and output spaces). Gridded (volumetric) resamplings were performed using antsApplyTransforms (ANTs), configured with Lanczos interpolation to minimize the smoothing effects of other kernels (Lanczos 1964). Non-gridded (surface) resamplings were performed using mri_vol2surf (FreeSurfer).

Many internal operations of *fMRIPrep* use *Nilearn* 0.8.1^21^ (RRID:SCR_001362), mostly within the functional processing workflow. For more details of the pipeline, see [the section corresponding to workflows in fMRIPrep’s documentation](https://fmriprep.readthedocs.io/en/latest/workflows.html).

##### S1.2.3 Copyright Waiver

The above boilerplate text was automatically generated by fMRIPrep with the express intention that users should copy and paste this text into their manuscripts *unchanged*. It is released under the [CC0](https://creativecommons.org/publicdomain/zero/1.0/) license.

#### S1.3 Region of Interest Selection

For resting-state fMRI data, ROIs were selected based on previous literature^22–24^, which included the medial prefrontal cortex (MPFC [3,54,-2]), PCC [0,-52,26], bilateral hippocampi (left [-29,-18,-16], right [29,-18,-16]), bilateral caudate (left [-10,14,0], right [10,14,0]), and bilateral putamen (left [-28,2,0], right [-28,2,0]). Masks were chosen using the WFU PickAtlas^25^ for the PCC, hippocampi, caudate, and putamen, and Stanford Willard Atlas^26^ for the MPFC. Masks further constrained the sphere around the MNI centroid.

#### S1.4 Quality Assurance Procedures

For resting-state data, quality assurance procedures included assessment of outliers (± 2 SD away from the mean) in framewise displacement, DVARS (the rate of change of BOLD signal across the entire brain at each frame of data), temporal signal-to-noise ratio, entropy focus criterion, high full width at half maximum (excessive image smoothness), signal-to-noise ratio, contrast-to-noise ratio, white matter/gray matter contrast, and global signal outliers. Following pre-processing, 8 sessions were removed as fMRIPrep revealed framewise displacement was excessive (>.50mm) and the percentage of movement for the total scan for those participants was greater than 20%. In total, 4 baseline, and 4 24-month follow-up sessions were removed. We also removed participants that had failed QC by an external team who provided the dataset, resulting in a further two baseline and three 24-month follow-up participants being removed.

For structural data, no subjects fell below the quality threshold of image quality rating (IQR) < 70%, and mean IQR was 85.18% (*SD =* 0.46 *range* = 84.50-86.03%). Quality assurance procedures included homogeneity checks across all structural scans, assessment of inter-scan similarity using correlation matrices, visual inspection of boxplots of mean correlation values to identify global outliers, and visual inspection of tissue segmentation maps (gray matter, white matter, CSF) for the 10 subjects with lowest quality metrics. Following pre-processing, outliers (± 2 SD away from the mean) with inconsistent quality across longitudinal sessions, extreme TIV values consistently across and with abnormal tissue proportions (*n* = 12) were assessed. Visual inspection of original T1-weighted images and segmentation overlays, assessment of segmentation boundary accuracy, cross-session comparison for longitudinal consistency and examination of tissue classification quality was performed. Of the 12 priority participants (38 sessions) identified by automated quality metrics as outliers, 1 participant (3 sessions) was excluded due to consistent motion artifacts and segmentation issues. For participants with motion artifacts but visible disease-related features (enlarged ventricles, atrophy), a more inclusive approach was adopted to avoid selection bias against more severely affected individuals.

#### S1.5 Linear Mixed Modelling

Intraclass correlation coefficients (ICC) quantified between-subject variability, substantiating the inclusion of subject-specific random effects. Restricted maximum likelihood estimation (REML) was used to obtain unbiased variance estimates by accounting for fixed effects degrees of freedom^27^. This approach accurately captured between-subject variability given the model structure (11 fixed effects) and sample-to-parameter ratio (n-p =22.1)^28^. Linear and quadratic session effects were compared to examine changes in depression symptoms over time. Model selection involved likelihood ratio tests (maximum likelihood estimation for nested models), information criteria evaluation (AIC and BIC) with preference for parsimonious solutions^29^, and marginal R² values quantified variance explained by fixed effects. Analyses were conducted in R version 4.1.3^30^ using lme4^31^ and lmerTest^31^.

#### S1.6 Dynamic Causal Modelling

Dynamic causal modelling (DCM), a popular method to infer effective connectivity, is a series of modelling techniques that are biophysically informed effective connectivity analyses of distributed neural networks^32^ .DCM broadly refers to the Bayesian modelling procedure, which allows the testing of each potential model and its corresponding hypothesis regarding the construction of the network^32,33^. In this way, it is a hypothesis-driven approach to understanding the neural networks underpinning observed brain activation ^33^. Model formulation evaluates how data are caused in the network, and DCM uses parameterized connections to understand the dynamics of intrinsic hidden states in the sources of fMRI data^32^. Model inversion allows neuronal states to influence other neuronal states^32^. DCM facilitates a mechanistic approach to understanding distributed neural processing and disturbances in connectivity changes in a pre-specified network of brain regions^32,33^. There are two key forms of DCM used for resting-state fMRI: stochastic DCM and spectral DCM.

To model resting-state activity, which occurs in the absence of external stimuli, we incorporate a stochastic component to account for neural fluctuations within the model. Mathematically, we express the formulation of the stochastic generative model through two equations. The first equation represents the neuronal state dynamics as follows:

$$x˙\left( t \right)=f\left( x\left( t \right),u\left( t \right),\theta\right)+v\left( t \right) (1)$$

The second equation represents the observation process, which is a static nonlinear mapping from the hidden physiological states in equation (1) to the observed BOLD activity:

$$y\left( t \right)=h\left( x\left( t \right),\phi\right)+e\left( t \right) (2)$$

Here, *x*˙(*t*) represents the rate of change of the neuronal states *x*(*t*), where *θ* denotes the unknown parameters, specifically the effective connectivity. The terms $v\left( t \right)$ and $e\left( t \right)$ correspond to stochastic processes referred to as state noise and measurement (or observation) noise, respectively. These processes model the random neuronal fluctuations responsible for driving the resting-state activity.

In equation (2), $\phi$ represents the unknown parameters associated with the hemodynamic observation function. Additionally, $u\left( t \right)$ signifies any exogenous (or experimental) inputs that influence the hidden states. It's important to note that such inputs are typically absent in resting-state designs.

Criticisms of using stochastic DCM for fMRI data include its unstable model inversion and high computational cost by accounting for neuronal variation in the time domain^24,34^. The use of spectral DCM ensures stability in estimations and computational efficiency^24,34^. Research has demonstrated that spectral DCM provides more accurate estimates that are sensitive to group differences, when compared to stochastic DCM^24^.

Spectral DCM introduces a constrained inversion approach to the stochastic model by parameterizing the neuronal fluctuations $v\left( t \right)$. This methodology simplifies the generative model by substituting the original time series with their second-order statistics, specifically the cross spectra. This implies that instead of estimating time-varying hidden states, we are now focused on estimating their time-invariant covariance. To achieve this, we need to estimate the covariance of the random fluctuations. In this context, a scale-free (power-law) form for the state noise (or observation noise) is employed:

$$g_{v}\left( \omega,\theta\right)=\alpha_{v}\omega{}^{-\beta v}$$

$$g_{e}\left( \omega,\theta\right)=\alpha_{e}\omega^{-\beta e} (3)$$

In these equations, the parameters $\{\alpha,\beta\}\subset\theta$ govern the amplitudes and exponents of the spectral density of the neural fluctuations. Notably, the parameterization of endogenous fluctuations results in deterministic states, simplifying the inversion scheme significantly. Consequently, the estimation process now focuses solely on the model's parameters and hyperparameters.

##### S1.6.1 Parametric Empirical Bayes

Parametric Empirical Bayes refers to the Bayesian inversion or fitting of models. In longitudinal models, constraints on the posterior density over model parameters at any given level are determined by the level above, and these constraints are referred to as empirical priors as they are informed by empirical data. The parametric empirical Bayes (PEB) model for DCM parameters elucidates how individual (within-subject) connections are derived from group membership. Parametric random effects modelling utilizes the complete posterior density over the parameters from each participant's DCM, encompassing both the expected strength of each connection and the associated uncertainty (i.e., posterior covariance), to inform the group-level result (i.e., group differences).

Mathematically, for DCM studies with *N* participants and *M* parameters per DCM, the responses of the *i*-th participant and the distribution of the parameters over participants can be modelled as:

$y_{i}=\Gamma_{i}^{\left( 1 \right)}\left( \theta^{\left( 1 \right)} \right)+ \varepsilon_{i}^{\left( 1 \right)}$

$\theta^{\left( 1 \right)}=\Gamma^{\left( 2 \right)}\left( \theta^{\left( 2 \right)} \right)+ \varepsilon^{\left( 2 \right)}$	(4)

$\theta^{\left( 2 \right)}=\eta+ \varepsilon^{\left( 3 \right)}$

Here, $y_{i}$ represents the BOLD time series from the *i*-th participant, and $\Gamma_{i}^{\left( 1 \right)}$ is a nonlinear mapping from the parameters of a model to the predicted response y, which in this study corresponds to the model in Eq. S1 above. $\varepsilon_{i}^{\left( 1 \right)}$represents independent and identically distributed observation noise (equivalent to $e\left( t \right)$ in Eq. S2).

$\Gamma^{\left( 2 \right)}\left( \theta^{\left( 2 \right)} \right)=(X\bigotimes W)\beta$	(5)

Here, $\beta\subset\theta$ represents group means or effects encoded by a design matrix comprising both between-subject $(X)$ and within-subject $\left( W \right)$components. The between-subject part encodes differences among subjects or covariates such as age, while the within-subject part specifies mixtures of parameters that exhibit random effects. It is assumed that the first column of the design matrix is a constant term that models group means, and subsequent columns encode group differences.

##### S1.6.2 Self-Connectivity in Dynamic Causal Modelling

Within the DCM framework, self-connections are modelled as inhibitory to prevent potential runaway excitation; however, these self-connection parameters are logarithmically scaled (using the transformation log(−2*a*) in SPM). This logarithmic scaling is employed to enhance the numerical stability of the model fitting procedures and is technically motivated by using log-normal priors to enforce recurrent self-inhibition. Consequently, these self-connections can take both positive and negative values, with specific interpretations.

In SPM's reporting convention, a zero value (arbitrarily set) for a self-connection corresponds to −0.5 Hz, representing the default prior self-connectivity value. A positive self-connection signifies a relative increase in inhibition (faster decay rates, below 0.5 Hz), while a negative self-connection indicates a relative decrease in inhibition (slower decay rates in the −0.5 to 0 Hz range). These inhibitory self-connections regulate the gain or sensitivity to inputs from other regions. Decreased self-inhibition implies increased synaptic gain or sensitivity to inputs, whereas increased self-inhibition suggests a reduction in synaptic gain or sensitivity to inputs. Importantly, only self-connections are subjected to this logarithmic scaling transformation in DCM.

#### S1.7 Assumptions

##### S1.7.1 Demographic Comparison Assumptions

Reported below are participant characteristics that exhibited violations of statistical assumptions for between-group comparisons. As these violations were confined to demographic variables rather than primary outcome measures, these assumption violations were deemed acceptable for the scope of this analysis.

Residuals for CAG repeat were not normally distributed (W = 0.946, *p* < 0.001), and the Q-Q plot showed residuals deviating at the tails. The box plot showed some outliers that contributed to non-normality and evidence of heteroscedasticity (eFigure 2).

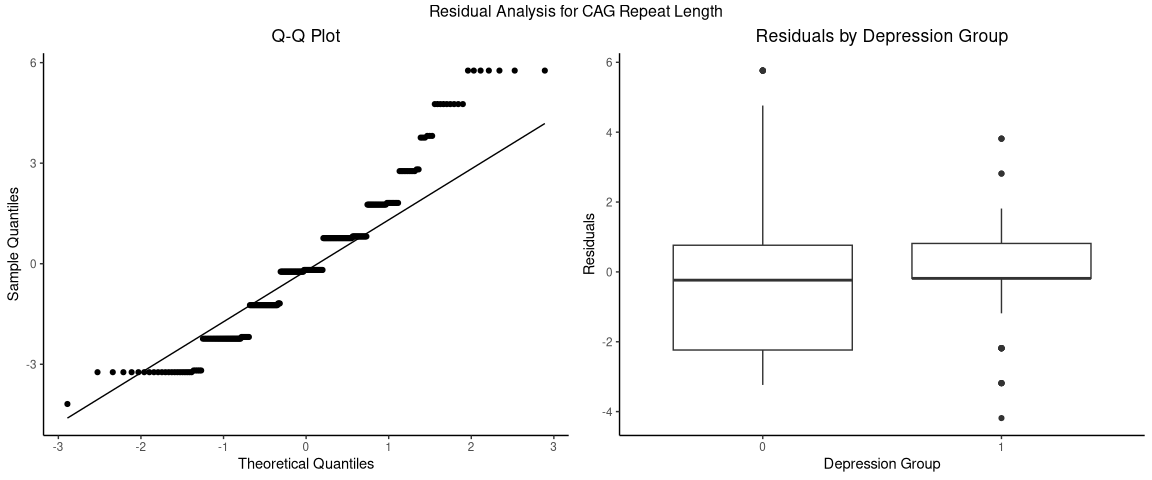

eFigure 2 Q-Q plot and boxplot of Residuals for Cytosine, Adenine, Guanine (CAG) Repeat Length. The left plot shows the results of the Q-Q plot of residuals for participant CAG repeat length. The right plot shows the results of the box plots of residuals for CAG repeat length, split by participants with and without a history of depression.

Residuals for DBS were normally distributed (W = 0.994, *p* = 0.466), as supported by visual inspection of the Q-Q plot. The box plot showed little evidence of heteroscedasticity (eFigure 3).

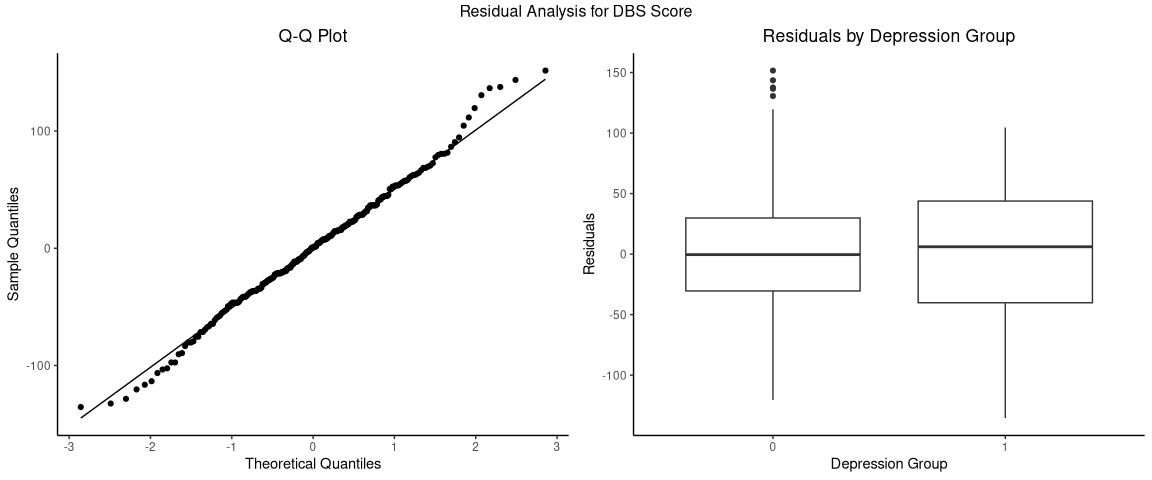

eFigure 3 Q-Q plot and boxplot of Residuals for Disease Burden Score (DBS). The left plot shows the results of the Q-Q plot of residuals for participant DBS. The right plot shows the results of the box plots of residuals for DBS, split by participants with and without a history of depression.

Residuals for BDI-II total score deviated from normality (W = 0.826, *p* <0.001), as supported by visual inspection of the Q-Q plot. The box plot showed some outliers that contributed to non-normality but little evidence of heteroscedasticity (eFigure 4).

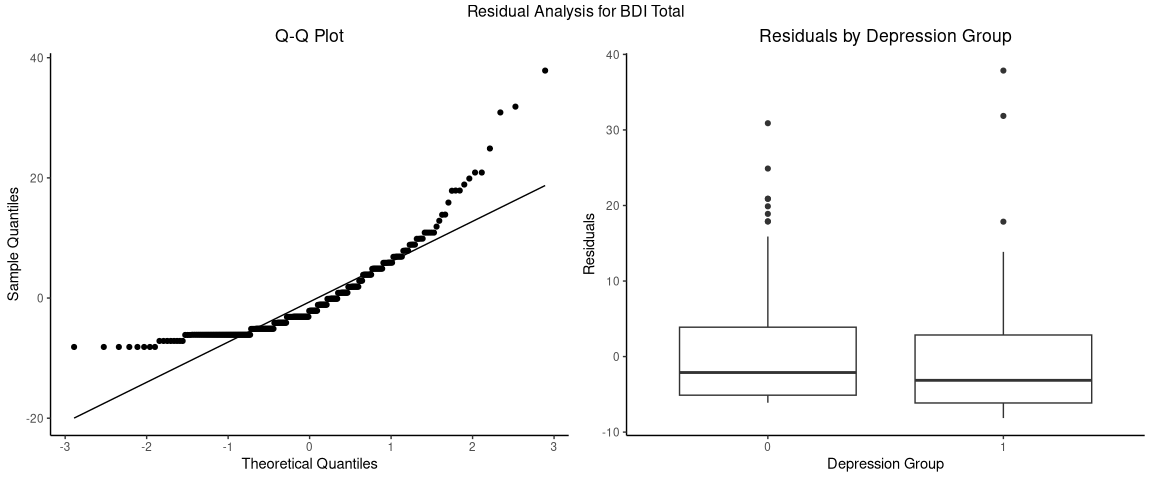

eFigure 4 Q-Q plot and boxplot of Residuals for Beck Depression Inventory, 2nd Edition (BDI-II) Score. The left plot shows the results of the Q-Q plot of residuals for participant BDI-II score. The right plot shows the results of the box plots of residuals for BDI-II score, split by participants with and without a history of depression.

Residuals for HADS-D score deviated from normality (W = 0.812, *p* <0.001) and the Q-Q plot showed residuals deviating at the tails. The box plot showed some outliers that contributed to non-normality but no evidence of heteroscedasticity (eFigure 5).

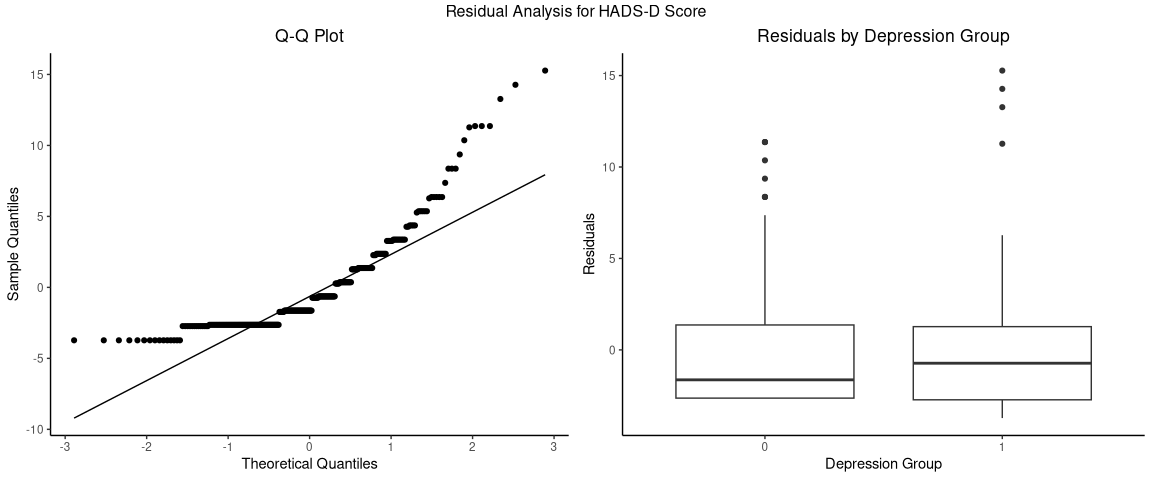

eFigure 5 Q-Q plot and boxplot of Residuals for Hospital Anxiety and Depression Scale, Depression Subscale (HADS-D) Score. The left plot shows the results of the Q-Q plot of residuals for participant HADS-D score. The right plot shows the results of the box plots of residuals for HADS-D score, split by participants with and without a history of depression.

##### S1.7.2 Linear Mixed Model Assumptions

Assumptions were met for both the BDI-II and HADS-D models. Visual inspection of Q-Q plots demonstrated acceptable normality of residuals despite minor deviations at extreme values, particularly for the BDI-II model (eFigure 6).

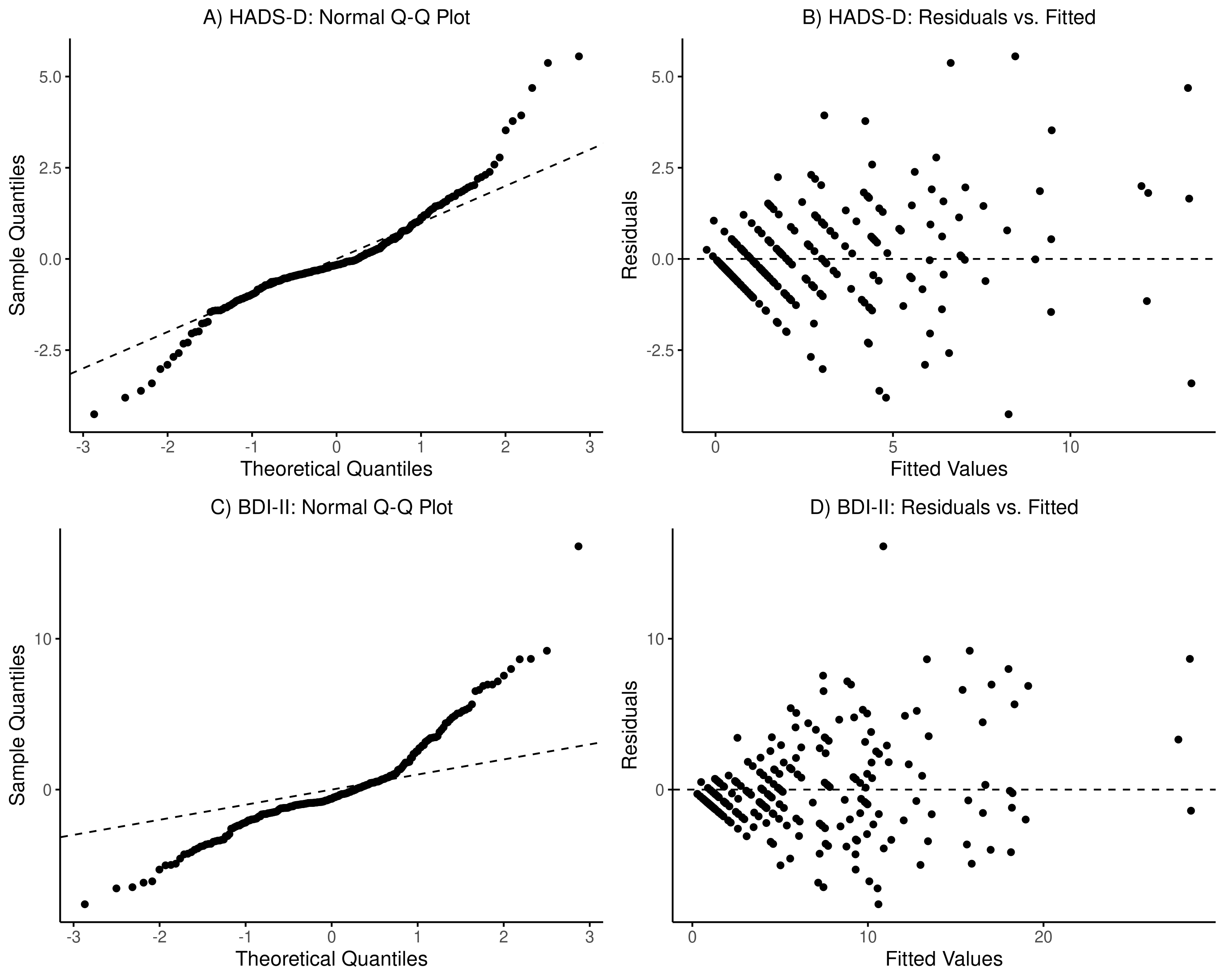

eFigure 6 Diagnostic Plots for Model Residuals. (a & c) Normal Q-Q plots for the HADS-D and BDI-II models respectively, showing generally linear alignment along the reference line with minor deviations at distribution extremes. (b & d) Residuals against fitted values, demonstrating appropriate random scatter without systematic patterns, confirming homoscedasticity assumptions.

Residual versus fitted value plots showed consistent variance across the prediction range demonstrating homoscedasticity. Random effects assumptions were satisfied, with Q-Q plots for subject-specific effects displaying appropriate normality and high intraclass correlation coefficients justifying hierarchical structure (eFigure 7).

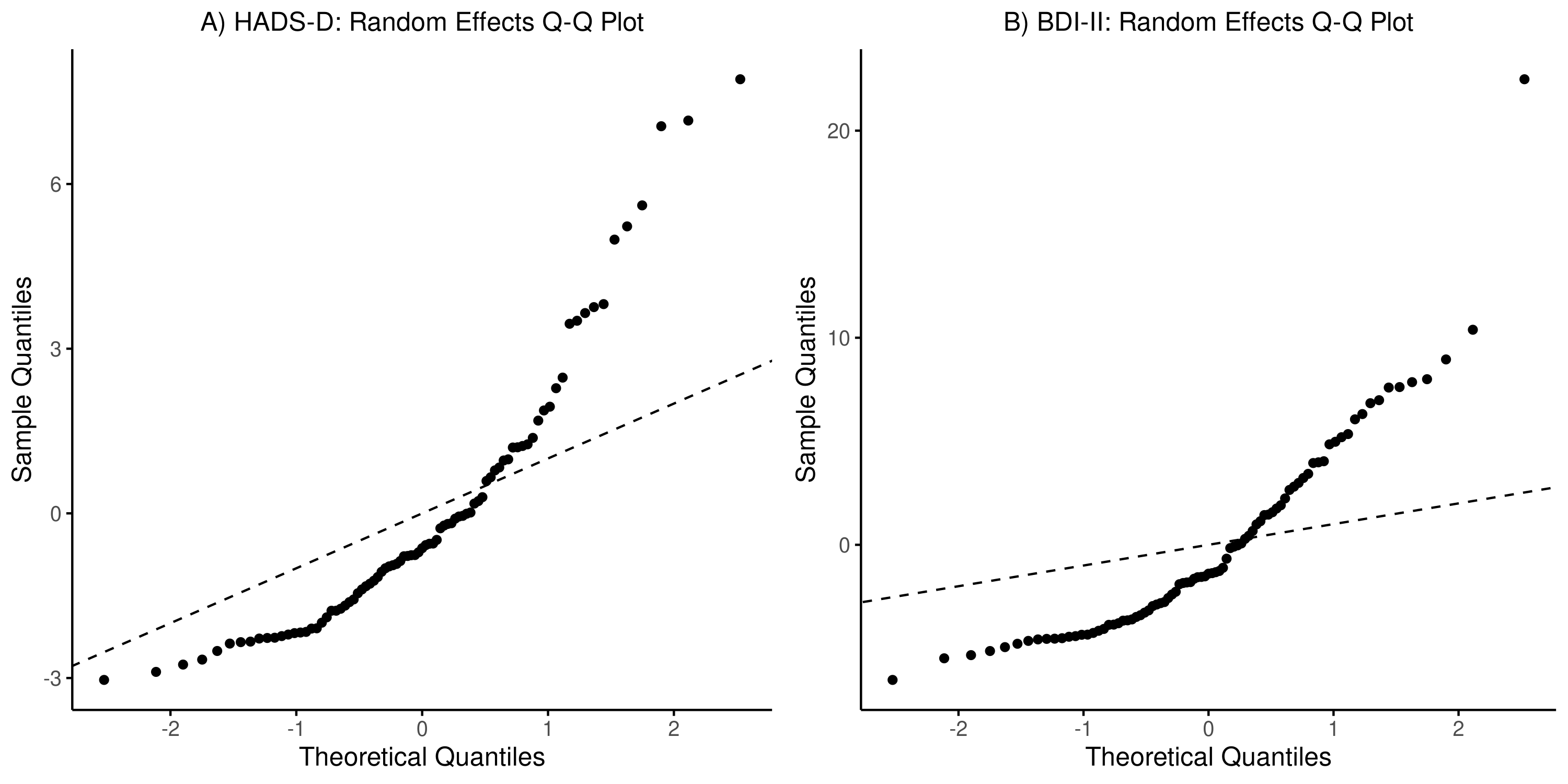

eFigure 7 Random Effects Normality Assessment. Q-Q plots of subject-specific random intercepts demonstrate acceptable normality in both (A) HADS-D and (B) BDI-II models.

Likelihood ratio tests comparing linear and quadratic temporal specifications (χ²(1) = 2.786, p = .095; BDI-II: χ²(1) = 0.358, p = .550), confirmed appropriateness of linear parameterization.

### S2. eResults

#### S2.1 Participant Characteristics

eTable 1 Depression History Classification Details

| **Variable** | **Baseline** | | | | **12 Months** | **24 Months** |
| --- | --- | --- | --- | --- | --- | --- |
| **Depression Status** |  | | | |  |  |
| Current episode | 16 (15.4%) | | | | 13 (15.3%) | 8 (11.0%) |
| Median duration (IQR) years | 4.0 (2.3-6.7) | | | | 4.9 (3.0-7.5) | 6.5 (5.6-8.5) |
| In remission | 15 (14.4%) | | | | 10 (11.8%) | 10 (13.7%) |
| Never depressed | 73 (70.2%) | | | | 62 (72.9%) | 55 (75.3%) |
| **Transitions Between Timepoints** |  | | | |  |  |
| Remission to new episode | - | | | | 1 (1.2%) | 0 (0.0%) |
| Episode to remission | - | | | | 0 (0.0%) | 1 (1.4%) |
| Stable episode | - | | | | 12 (14.3%) | 8 (11.4%) |
| Stable remission | - | | | | 10 (11.9%) | 8 (11.4%) |
| **ICD-10 Diagnostic Category (all timepoints)** | |  |  |  | | |
| F32.9 (Depressive Episode, Unspecified) | | 66 (91.7%) | | | | |
| F32.0 (Mild Depressive Episode) | | 3 (4.2%) | | | | |
| F32.2 (Severe Depressive Episode without Psychotic Symptoms) | | 3 (4.2%) | | | | |

ICD = International Statistical Classification of Diseases and Related Health Problems 10th Revision

eTable 2 Mood Medication Use Details at Baseline

| **Medication** | **Total** | | **Depression** | | **No Depression** | | **Duration (years)  Mean (SD), Range** | | **Medication Dose (mg) Mean (SD), Range** | | **Listed Indication** | | **Regime** |
| --- | --- | --- | --- | --- | --- | --- | --- | --- | --- | --- | --- | --- | --- |
| **Antipsychotics** |  | |  | |  | |  | |  | |  | |  |
| Aripiprazole | 1 | | 1 | | 0 | | 0.31 (0.00), 0.31-0.31 | | 15.0 (0.0), 15.0-15.0 | | Depression (100%) | | once daily (100%) |
| Olanzapine | 1 | | 1 | | 0 | | 2.06 (0.00), 2.06-2.06 | | 2.5 (0.0), 2.5-2.5 | | Irritability (100%) | | once daily (100%) |
| Quetiapine | 2 | | 2 | | 0 | | 1.37 (1.23), 0.50-2.24 | | 75.0 (35.4), 50.0-100.0 | | Depression (100%) | | once daily (100%) |
| **Benzodiazepines** | |  | |  | |  | |  | |  | |  | |
| Bromazepam | 1 | | 0 | | 1 | | 3.24 (0.00), 3.24-3.24 | | 1.5 (0.0), 1.5-1.5 | | Anxiety (100%) | | once daily (100%) |
| Oxazepam | 1 | | 0 | | 1 | | 0.41 (0.00), 0.41-0.41 | | 10.0 (0.0), 10.0-10.0 | | Anxiety (100%) | | twice daily (100%) |
| Temazepam | 1 | | 1 | | 0 | | 14.83 (0.00), 14.83-14.83 | | 10.0 (0.0), 10.0-10.0 | | Sleep disorder (100%) | | once daily (100%) |
| **Non-SSRI** |  | |  | |  | |  | |  | |  | |  |
| Amitriptyline | 1 | | 1 | | 0 | | 13.78 (0.00), 13.78-13.78 | | 50.0 (0.0), 50.0-50.0 | | Post-traumatic Stress (100%) | | once daily (100%) |
| Bupropion | 3 | | 2 | | 1 | | 1.59 (0.94), 0.66-2.89 | | 200.0 (70.7), 150.0-300.0 | | Anxiety (33%), Depression (67%) | | once daily (100%) |
| Desvenlafaxine | 1 | | 1 | | 0 | | 2.37 (0.00), 2.37-2.37 | | 100.0 (0.0), 100.0-100.0 | | Depression (100%) | | once daily (100%) |
| Duloxetine | 2 | | 2 | | 0 | | 1.73 (0.49), 1.23-2.22 | | 60.0 (0.0), 60.0-60.0 | | Depression (100%) | | once daily (100%) |
| Mianserin | 1 | | 0 | | 1 | | 0.41 (0.00), 0.41-0.41 | | 30.0 (0.0), 30.0-30.0 | | Depression (100%) | | once daily (100%) |
| Mirtazapine | 1 | | 1 | | 0 | | 1.04 (0.00), 1.04-1.04 | | 15.0 (0.0), 15.0-15.0 | | Depression (100%) | | once daily (100%) |
| Trazodone | 1 | | 1 | | 0 | | 0.51 (0.00), 0.51-0.51 | | 50.0 (0.0), 50.0-50.0 | | Depression (100%) | | once daily (100%) |
| Venlafaxine | 2 | | 2 | | 0 | | 0.74 (0.33), 0.51-0.98 | | 93.8 (26.5), 75.0-112.5 | | Depression (100%) | | once daily (50%), twice daily (50%) |
| **SSRI** |  | |  | |  | |  | |  | |  | |  |
| Citalopram | 12 | | 6 | | 6 | | 1.96 (1.68), 0.38-5.73 | | 20.8 (7.9), 10.0-40.0 | | Anxiety (25%), Depression (75%) | | once daily (100%) |
| Escitalopram | 4 | | 1 | | 3 | | 3.38 (3.65), 0.00-7.82 | | 13.8 (4.8), 10.0-20.0 | | Depression (75%), Irritability (25%) | | once daily (100%) |
| Fluoxetine | 1 | | 1 | | 0 | | 3.87 (0.00), 3.87-3.87 | | 20.0 (0.0), 20.0-20.0 | | Depression (100%) | | once daily (100%) |
| Paroxetine | 2 | | 1 | | 1 | | 9.42 (11.68), 1.16-17.68 | | 10.2 (13.8), 0.5-20.0 | | Anxiety (50%), Depression (50%) | | once daily (100%) |

SSRI = Selective Serotonin Reuptake Inhibitors. All doses in mg, unless otherwise stated.

#### S2.2 Volume-based Morphometry

##### S2.2.1 Participant Characteristics

eTable 3 VBM Longitudinal Participant Characteristics for Variables of Interest

|  | | | **Baseline** | | | | **24-month follow-up** | | |
| --- | --- | --- | --- | --- | --- | --- | --- | --- | --- |
|  | **Depression History (N = 20)** | | | **No Depression History (N = 51)** | | **Depression History (N = 20)** | | | **No Depression History (N = 51)** |
| **Depression Status** |  | | |  | |  | | |  |
| Current episode | 11 (55.0%) | | | 0 (0.0%) | | 10 (50.0%) | | | 0 (0.0%) |
| Median duration (IQR) years | 4.0 (0.6-6.1) | | | N/A | | 6.2 (2.2-8.4) | | | N/A |
| In remission | 9 (45.0%) | | | 0 (0.0%) | | 10 (50.0%) | | | 0 (0.0%) |
| Never depressed | 0 (0.0%) | | | 51 (100.0%) | | 0 (0.0%) | | | 51 (100.0%) |
| **ICD-10 Diagnostic Category** |  | | |  | |  | | |  |
| F32.9 (Depressive Episode, Unspecified) | 18 (90.0%) | | | - | | 18 (90.0%) | | | - |
| F32.0 (Mild Depressive Episode) | 1 (5.0%) | | | - | | 1 (5.0%) | | | - |
| F32.2 (Severe Depressive Episode without Psychotic Symptoms) | 1 (5.0%) | | | - | | 1 (5.0%) | | | - |
| **Medication** |  | | |  | |  | | |  |
| Antipsychotics | 3 (15.0%) | | | 0 (0.0%) | | 4 (20.0%) | | | 0 (0.0%) |
| Benzodiazepines | 1 (5.0%) | | | 2 (3.9%) | | 2 (10.0%) | | | 1 (2.0%) |
| SSRI | 5 (25.0%) | | | 5 (9.8%) | | 6 (30.0%) | | | 7 (13.7%) |
| Number of Medications - Mean(SD), range | 0.6 (0.9), 0-3 | | | 0.2 (0.5), 0-3 | | 0.8 (1.0), 0-3 | | | 0.2 (0.4), 0-2 |
| **HD-related Variables** | |  | | |  | | |  | |
| HD-ISS Stage |  | | |  | |  | | |  |
| 0 | 3 (15.0%) | | | 20 (39.2%) | | 3 (15.0%) | | | 13 (25.5%) |
| 1 | 12 (60.0%) | | | 20 (39.2%) | | 7 (35.0%) | | | 23 (45.1%) |
| 2 | 5 (25.0%) | | | 11 (21.6%) | | 10 (50.0%) | | | 14 (27.5%) |
| 3 | 0 (0.0%) | | | 0 (0.0%) | | 0 (0.0%) | | | 1 (2.0%) |
| **Depression Variables** | |  | | |  | | |  | |
| Current Depression |  | | |  | |  | | |  |
| No Current Depression | 9 (45.0%) | | | 51 (100.0%) | | 10 (50.0%) | | | 51 (100.0%) |
| Current Depression | 11 (55.0%) | | | 0 (0.0%) | | 10 (50.0%) | | | 0 (0.0%) |
| BDI-II Score |  | | |  | |  | | |  |
| Mean (SD) | 8.3 (7.6) | | | 5.3 (6.9) | | 7.5 (6.4) | | | 5.5 (5.9) |
| Range | 0-26 | | | 0-27 | | 0-21 | | | 0-31 |
| HADS-D Score |  | | |  | |  | | |  |
| Mean (SD) | 4.0 (4.1) | | | 2.4 (3.5) | | 3.4 (3.6) | | | 2.6 (3.2) |
| Range | 0-18 | | | 0-14 | | 0-15 | | | 0-14 |
| BDI-II Clinical Cut-off |  | | |  | |  | | |  |
| Clinically Sig. Symptoms | 5 (25.0%) | | | 10 (19.6%) | | 4 (20.0%) | | | 10 (19.6%) |
| Low Depressive Symptoms | 15 (75.0%) | | | 41 (80.4%) | | 16 (80.0%) | | | 41 (80.4%) |
| HADS-D Clinical Cut-off | |  | | |  | | |  | |
| Clinically Sig. Symptoms | 4 (20.0%) | | | 6 (11.8%) | | 3 (15.0%) | | | 5 (9.8%) |
| Low Depressive Symptoms | 16 (80.0%) | | | 45 (88.2%) | | 17 (85.0%) | | | 46 (90.2%) |

IQR = Interquartile Range; ICD-10 = International Statistical Classification of Diseases and Related Health Problems 10th Revision; SSRI = Selective Serotonin Reuptake Inhibitors; HD-ISS = Huntington’s Disease Integrated Staging System; BDI-II = Beck Depression Inventory-II; HADS-D = Beck Depression Inventory-II.

#### S2.3 Linear Mixed Modelling

##### S2.3.1 Participant Characteristics

eTable 4 LMM Longitudinal Participant Characteristics for Variables of Interest

|  | **Baseline** | | | | | **12-month follow-up** | | | | **24-month follow-up** | | |
| --- | --- | --- | --- | --- | --- | --- | --- | --- | --- | --- | --- | --- |
|  | **Depression History (N = 26)** | | | | **No Depression History (N = 60)** | **Depression History (N = 25)** | | **No Depression History (N = 60)** | | **Depression History (N = 20)** | | **No Depression History (N = 52)** |
| **Depression Status** |  | | | |  |  | |  | |  | |  |
| Current episode | 14 (53.8%) | | | | 0 (0.0%) | 15 (60.0%) | | 0 (0.0%) | | 10 (50.0%) | | 0 (0.0%) |
| Median duration (IQR) years | 4.0 (1.5-6.5) | | | | N/A | 4.9 (1.5-7.0) | | N/A | | 6.2 (2.2-8.4) | | N/A |
| In remission | 12 (46.2%) | | | | 0 (0.0%) | 10 (40.0%) | | 0 (0.0%) | | 10 (50.0%) | | 0 (0.0%) |
| Never depressed | 0 (0.0%) | | | | 60 (100.0%) | 0 (0.0%) | | 60 (100.0%) | | 0 (0.0%) | | 52 (100.0%) |
| **ICD-10 Diagnostic Category** | | |  | |  |  | |  | |  | |  |
| F32.9 (Depressive Episode, Unspecified) | 24 (92.3%) | | | | 0 (0.0%) | 23 (92.0%) | | 0 (0.0%) | | 18 (90.0%) | | 0 (0.0%) |
| F32.0 (Mild Depressive Episode) | 1 (3.8%) | | | | 0 (0.0%) | 1 (4.0%) | | 0 (0.0%) | | 1 (5.0%) | | 0 (0.0%) |
| F32.2 (Severe Depressive Episode without Psychotic Symptoms) | 1 (3.8%) | | | | 0 (0.0%) | 1 (4.0%) | | 0 (0.0%) | | 1 (5.0%) | | 0 (0.0%) |
| **Medication** |  | | | |  |  | |  | |  | |  |
| Antipsychotics | 3 (11.5%) | | | | 0 (0.0%) | 3 (12.0%) | | 0 (0.0%) | | 4 (20.0%) | | 0 (0.0%) |
| Benzodiazepines | 1 (3.8%) | | | | 2 (3.3%) | 2 (8.0%) | | 1 (1.7%) | | 2 (10.0%) | | 1 (1.9%) |
| SSRI | 5 (19.2%) | | | | 7 (11.7%) | 5 (20.0%) | | 10 (16.7%) | | 6 (30.0%) | | 7 (13.5%) |
| Number of Medications - Mean(SD), range | 0.6 (0.8), 0-3 | | | | 0.2 (0.5), 0-3 | 0.7 (0.8), 0-3 | | 0.2 (0.6), 0-3 | | 0.8 (1.0), 0-3 | | 0.2 (0.4), 0-2 |
| **HD-related Variables** | |  | |  | | |  | |  | |  | |
| HD-ISS Stage |  | | | |  |  | |  | |  | |  |
| 0 | 4 (15.4%) | | | | 22 (36.7%) | 3 (12.0%) | | 18 (30.0%) | | 3 (15.0%) | | 13 (25.0%) |
| 1 | 14 (53.8%) | | | | 25 (41.7%) | 14 (56.0%) | | 23 (38.3%) | | 7 (35.0%) | | 24 (46.2%) |
| 2 | 7 (26.9%) | | | | 13 (21.7%) | 7 (28.0%) | | 16 (26.7%) | | 10 (50.0%) | | 14 (26.9%) |
| 3 | 1 (3.8%) | | | | 0 (0.0%) | 1 (4.0%) | | 3 (5.0%) | | 0 (0.0%) | | 1 (1.9%) |
| **Depression Variables** | |  | |  | | |  | |  | |  | |
| Current Depression | |  | |  | | |  | |  | |  | |
| No Depression | 12 (46.2%) | | | | 60 (100.0%) | 10 (40.0%) | | 60 (100.0%) | | 10 (50.0%) | | 52 (100.0%) |
| Current Depression | 14 (53.8%) | | | | 0 (0.0%) | 15 (60.0%) | | 0 (0.0%) | | 10 (50.0%) | | 0 (0.0%) |
| BDI-II Score |  | | | |  |  | |  | |  | |  |
| Mean (SD) | 7.9 (7.0) | | | | 5.7 (7.2) | 8.4 (6.2) | | 6.0 (7.1) | | 7.5 (6.4) | | 5.4 (5.8) |
| Range | 0-26 | | | | 0-27 | 0-22 | | 0-37 | | 0-21 | | 0-31 |
| HADS-D Score |  | | | |  |  | |  | |  | |  |
| Mean (SD) | 3.8 (3.7) | | | | 2.6 (3.6) | 3.3 (3.0) | | 2.3 (3.0) | | 3.4 (3.6) | | 2.5 (3.1) |
| Range | 0-18 | | | | 0-14 | 0-10 | | 0-13 | | 0-15 | | 0-14 |
| BDI-II Clinical Cut-off | |  | |  | | |  | |  | |  | |
| Clinically Sig. Symptoms | 6 (23.1%) | | | | 13 (21.7%) | 9 (36.0%) | | 13 (21.7%) | | 4 (20.0%) | | 10 (19.2%) |
| Low Depressive Symptoms | 20 (76.9%) | | | | 47 (78.3%) | 16 (64.0%) | | 47 (78.3%) | | 16 (80.0%) | | 42 (80.8%) |
| HADS-D Clinical Cut-off | |  | |  | | |  | |  | |  | |
| Clinically Sig. Symptoms | 5 (19.2%) | | | | 8 (13.3%) | 4 (16.0%) | | 6 (10.0%) | | 3 (15.0%) | | 5 (9.6%) |
| Low Depressive Symptoms | 21 (80.8%) | | | | 52 (86.7%) | 21 (84.0%) | | 54 (90.0%) | | 17 (85.0%) | | 47 (90.4%) |

IQR = Interquartile Range; ICD-10 = International Statistical Classification of Diseases and Related Health Problems 10th Revision; SSRI = Selective Serotonin Reuptake Inhibitors; HD-ISS = Huntington’s Disease Integrated Staging System; BDI-II = Beck Depression Inventory-II; HADS-D = Beck Depression Inventory-II.

##### S2.3.2 LMM Findings

eTable 5 ****Fixed Effects from Linear Mixed Models Predicting Depression Scores****

| **Predictor** | **HADS-D** | | **BDI-II** | |
| --- | --- | --- | --- | --- |
|  | **β (SE)** | **p** | **β (SE)** | **p** |
| **Time Effects** |  |  |  |  |
| Time (Linear) | -0.029 (0.141) | 0.839 | -0.009 (0.316) | 0.978 |
| **Clinical Variables** |  |  |  |  |
| HD-ISS Stage | -0.039 (0.252) | 0.878 | 0.365 (0.560) | 0.515 |
| Depression History | 0.949 (0.823) | 0.251 | 2.077 (1.715) | 0.229 |
| Mood Medication Use | 2.050 (0.674) | 0.003 | 3.928 (1.462) | 0.008 |
| Depression History × Medication Use | -1.159 (1.140) | 0.311 | -2.653 (2.454) | 0.281 |
| **Grey Matter Volumes** |  |  |  |  |
| Caudate Volume | -7.505 (10.742) | 0.486 | -10.160 (22.287) | 0.650 |
| Putamen Volume | 6.029 (5.905) | 0.309 | 13.350 (12.424) | 0.285 |
| Hippocampus Volume | -4.446 (9.842) | 0.652 | 2.078 (20.447) | 0.919 |
| PCC Volume | 12.242 (5.415) | 0.026 | 10.753 (11.202) | 0.340 |
| MPFC Volume | -7.328 (4.195) | 0.084 | -7.122 (8.705) | 0.415 |

Values are coefficient (standard error). *p <0.05. All brain volumes are normalized by total intracranial volume (TIV) and scaled (×100). PCC = Posterior Cingulate Cortex; MPFC = Medial Prefrontal Cortex.

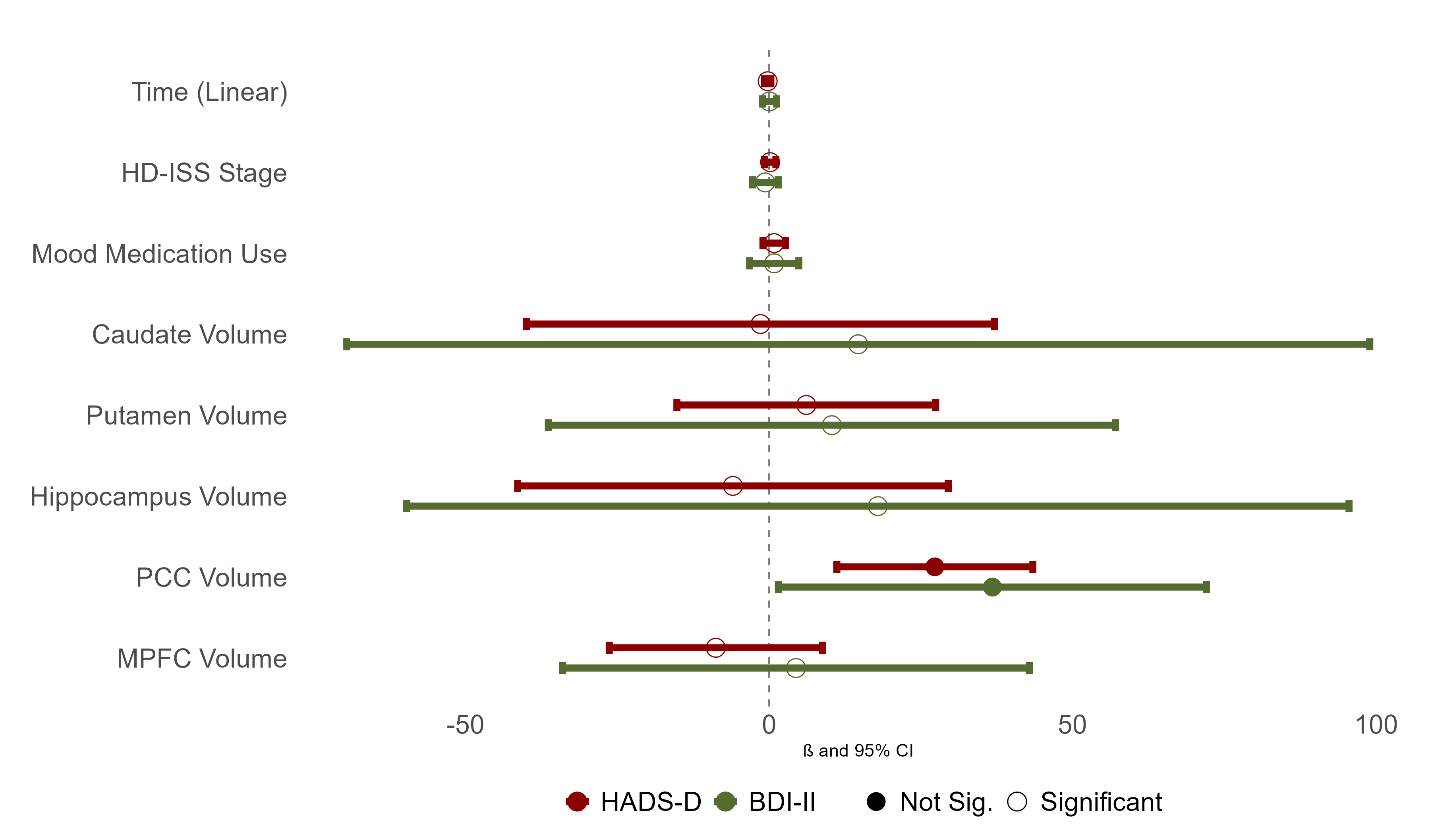

eFigure 8 ****Predictors of depressive symptomatology for HDGECs with a history of depression.** Forest plot displaying standardized β coefficients and 95% confidence intervals from linear mixed-effects models predicting Beck Depression Inventory-II (BDI-II; orange) and Hospital Anxiety and Depression Scale-Depression subscale (HADS-D; blue). Fixed effects include clinical variables (disease severity [HD-ISS], antidepressant use, history x medication interaction). All brain volumes are normalized by total intracranial volume (TIV) and scaled (×100) (caudate, putamen, hippocampus, posterior cingulate cortex [PCC], medial prefrontal cortex [MPFC]). Filled circles indicate statistical significance.**

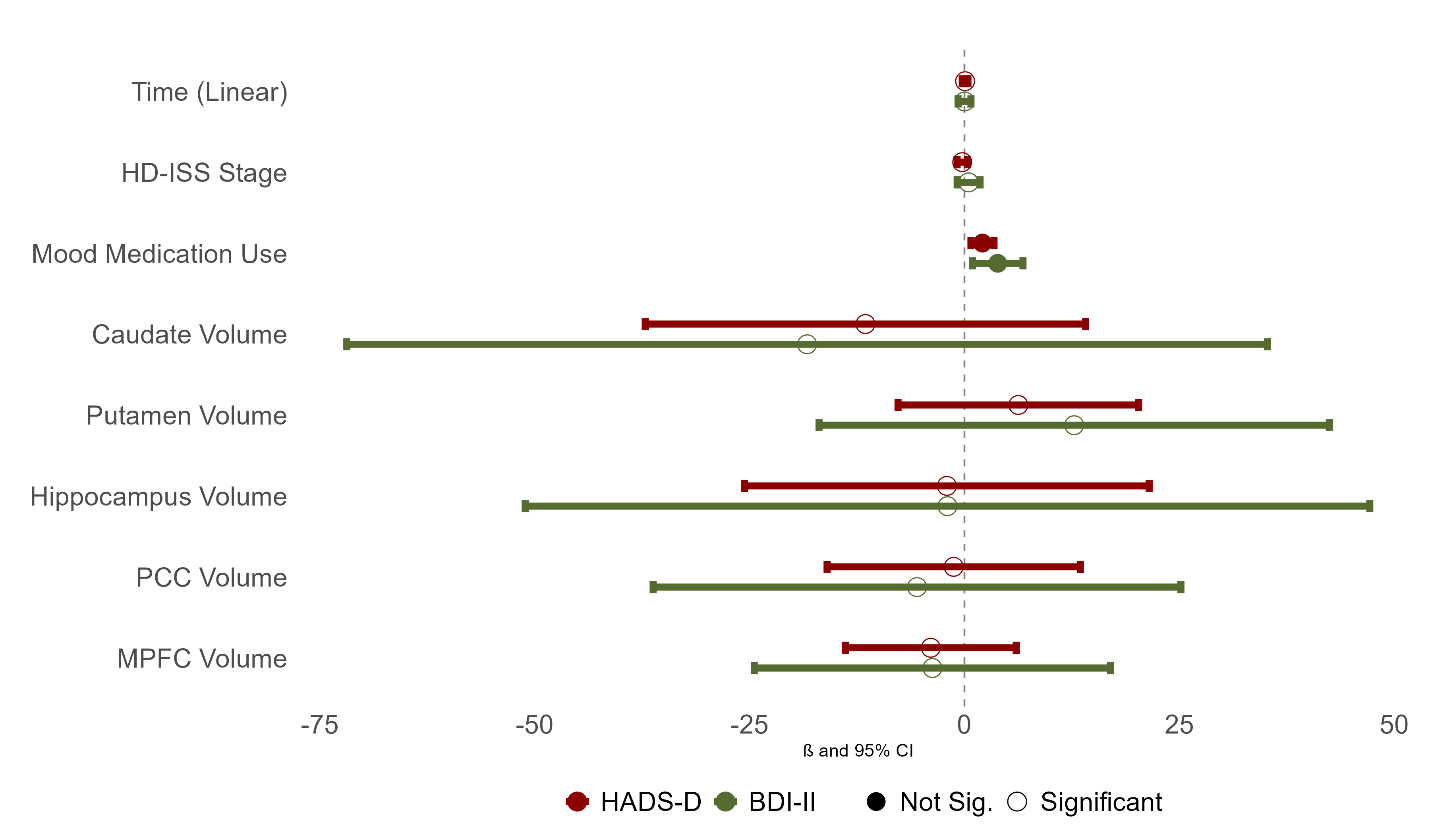

eFigure 9 ****Predictors of depressive symptomatology for HDGECs with no history of depression.** Forest plot displaying standardized β coefficients and 95% confidence intervals from linear mixed-effects models predicting Beck Depression Inventory-II (BDI-II; orange) and Hospital Anxiety and Depression Scale-Depression subscale (HADS-D; blue). Fixed effects include clinical variables (disease severity [HD-ISS], antidepressant use, history x medication interaction). All brain volumes are normalized by total intracranial volume (TIV) and scaled (×100) (caudate, putamen, hippocampus, posterior cingulate cortex [PCC], medial prefrontal cortex [MPFC]). Filled circles indicate statistical significance.**

#### S2.4 Spectral Dynamic Causal Modelling

##### S2.4.1 Participant Characteristics

eTable 6 DCM Longitudinal Participant Characteristics for Variables of Interest

|  | | **Baseline** | | | **24-month follow-up** | | |
| --- | --- | --- | --- | --- | --- | --- | --- |
|  | | **Depression History (N = 30)** | **No Depression History (N = 68)** | | **Depression History (N = 17)** | | **No Depression History (N = 49)** |
| **Depression Status** | |  |  | |  | |  |
| Current episode | | 18 (60.0%) | 0 (0.0%) | | 10 (58.8%) | | 0 (0.0%) |
| Median duration (IQR) years | | 3.9 (1.5-6.5) | N/A | | 6.2 (2.2-8.4) | | N/A |
| In remission | | 12 (40.0%) | 0 (0.0%) | | 7 (41.2%) | | 0 (0.0%) |
| Never depressed | | 0 (0.0%) | 68 (100.0%) | | 0 (0.0%) | | 49 (100.0%) |
| **ICD-10 Diagnostic Category** | |  |  | |  | |  |
| F32.9 (Depressive Episode, Unspecified) | | 29 (96.7%) | 0 (0.0%) | | 15 (88.2%) | | 0 (0.0%) |
| F32.0 (Mild Depressive Episode) | | 0 (0.0%) | 0 (0.0%) | | 1 (5.9%) | | 0 (0.0%) |
| F32.2 (Severe Depressive Episode without Psychotic Symptoms) | | 1 (3.3%) | 0 (0.0%) | | 1 (5.9%) | | 0 (0.0%) |
| **Medication** | |  |  | |  | |  |
| Antipsychotics | | 4 (13.3%) | 0 (0.0%) | | 4 (23.5%) | | 0 (0.0%) |
| Benzodiazepines | | 1 (3.3%) | 2 (2.9%) | | 1 (5.9%) | | 1 (2.0%) |
| SSRI | | 9 (30.0%) | 10 (14.7%) | | 6 (35.3%) | | 7 (14.3%) |
| Number of Medications - Mean(SD), range | | 0.8 (0.8), 0-3 | 0.2 (0.5), 0-3 | | 0.9 (1.0), 0-3 | | 0.2 (0.4), 0-2 |
| **HD-related Variables** |  | | |  | |  | |
| HD-ISS Stage | |  |  | |  | |  |
| 0 | | 5 (16.7%) | 21 (30.9%) | | 1 (5.9%) | | 12 (24.5%) |
| 1 | | 13 (43.3%) | 27 (39.7%) | | 6 (35.3%) | | 21 (42.9%) |
| 2 | | 8 (26.7%) | 18 (26.5%) | | 10 (58.8%) | | 14 (28.6%) |
| 3 | | 4 (13.3%) | 2 (2.9%) | | 0 (0.0%) | | 2 (4.1%) |
| **Depression Variables** |  | | |  | |  | |
| Current Depression | |  |  | |  | |  |
| No Current Depression | | 12 (40.0%) | 68 (100.0%) | | 7 (41.2%) | | 49 (100.0%) |
| Current Depression | | 18 (60.0%) | 0 (0.0%) | | 10 (58.8%) | | 0 (0.0%) |
| BDI-II Score | |  |  | |  | |  |
| Mean (SD) | | 9.6 (11.3) | 6.0 (7.2) | | 8.3 (6.6) | | 5.7 (6.0) |
| Range | | 0-46 | 0-27 | | 0-21 | | 0-31 |
| HADS-D Score | |  |  | |  | |  |
| Mean (SD) | | 4.7 (5.1) | 2.6 (3.5) | | 3.8 (3.8) | | 2.8 (3.3) |
| Range | | 0-19 | 0-14 | | 0-15 | | 0-14 |
| BDI-II Clinical Cut-off | |  |  | |  | |  |
| Clinically Sig. Symptoms | | 8 (26.7%) | 16 (23.5%) | | 4 (23.5%) | | 10 (20.4%) |
| Low Depressive Symptoms | | 22 (73.3%) | 52 (76.5%) | | 13 (76.5%) | | 39 (79.6%) |
| HADS-D Clinical Cut-off |  | | |  | |  | |
| Clinically Sig. Symptoms | | 7 (23.3%) | 9 (13.2%) | | 3 (17.6%) | | 6 (12.2%) |
| Low Depressive Symptoms | | 23 (76.7%) | 59 (86.8%) | | 14 (82.4%) | | 43 (87.8%) |

IQR = Interquartile Range; ICD-10 = International Statistical Classification of Diseases and Related Health Problems 10th Revision; SSRI = Selective Serotonin Reuptake Inhibitors; HD-ISS = Huntington’s Disease Integrated Staging System; BDI-II = Beck Depression Inventory-II; HADS-D = Beck Depression Inventory-II.

e25.Maldjian J, Laurienti P, Kraft R, Burdette J. An automated method for neuroanatomic and cytoarchitectonic atlas-based interrogation of fMRI data sets. *Neuroimage*. Published online July 25, 2003.

e26.Altmann A, Ng B, Landau SM, Jagust WJ, Greicius MD. Regional brain hypometabolism is unrelated to regional amyloid plaque burden. *Brain*. 2015;138(12):3734-3746. doi:10.1093/brain/awv278

e27.Tabachnick BG, Fidell LS. Multilevel Linear Modeling. In: *Using Multivariate Statistics*. Seventh edition. Always learning. Pearson; 2019:613-671.

e28.Molenberghs G, Verbeke G, eds. Estimation of the Marginal Model. In: *Linear Mixed Models for Longitudinal Data*. Springer New York; 2000:41-54. doi:10.1007/978-1-4419-0300-6_5

e29.Burnham KP, Anderson DR. Multimodel inference: understanding AIC and BIC in model selection. *Sociological methods & research*. 2004;33(2):261-304.

e30.R Core Team. *R: A Language and Environment for Statistical Computing*. R Foundation for Statistical Computing; 2022. https://www.R-project.org/

e31.Kuznetsova A, Brockhoff PB, Christensen RHB. lmerTest Package: Tests in Linear Mixed Effects Models. *J Stat Soft*. 2017;82(13):1-26. doi:10.18637/jss.v082.i13

e32.Friston KJ. Functional and Effective Connectivity: A Review. *Brain Connect*. 2011;1(1):13-36. doi:10.1089/brain.2011.0008

e33.Friston KJ, Harrison L, Penny W. Dynamic causal modelling. *NeuroImage*. 2003;19(4):1273-1302. doi:10.1016/S1053-8119(03)00202-7

e34.Friston KJ, Kahan J, Biswal B, Razi A. A DCM for resting state fMRI. *NeuroImage*. 2014;94:396-407. doi:10.1016/j.neuroimage.2013.12.009
